# BeeDNA: microfluidic environmental DNA metabarcoding as a tool for connecting plant and pollinator communities

**DOI:** 10.1101/2021.11.11.468290

**Authors:** Lynsey R. Harper, Matthew L. Niemiller, Joseph B. Benito, Lauren E. Paddock, E. Knittle, Brenda Molano-Flores, Mark A. Davis

## Abstract

Pollinators are imperiled by global declines that can reduce plant reproduction, erode essential ecosystem services and resilience, and drive economic losses. Monitoring pollinator biodiversity trends is key for adaptive conservation and management, but conventional surveys are often costly, time consuming, and require taxonomic expertise. Environmental DNA (eDNA) metabarcoding surveys are booming due to their rapidity, non-invasiveness, and cost efficiency. Microfluidic technology allows multiple primer sets from different markers to be used in eDNA metabarcoding for more comprehensive species inventories whilst minimizing biases associated with individual primer sets. We evaluated microfluidic eDNA metabarcoding for pollinator community monitoring by introducing a bumblebee colony to a greenhouse flower assemblage and sampling natural flower plots. We collected nectar draws, flower swabs, or whole flower heads from four flowering species, including two occurring in both the greenhouse and field. Samples were processed using two eDNA isolation protocols before amplification with 15 primer sets for two markers (COI and 16S). Microfluidic eDNA metabarcoding detected the target bumblebee and greenhouse insects as well as common regional arthropods. Pollinator detection was maximized using whole flower heads preserved in ATL buffer and extracted with a modified Qiagen^®^ DNeasy protocol for amplification with COI primers. eDNA surveillance could enhance pollinator assessment by detecting protected and endangered species and being more applicable to remote, inaccessible locations, whilst reducing survey time, effort, and expense. Microfluidic eDNA metabarcoding requires optimization to address remaining efficacy concerns but this approach shows potential in revealing complex networks underpinning critical ecosystem functions and services, enabling more accurate assessments of ecosystem resilience.

## 1. Introduction

Biodiversity is being lost at an alarming rate across habitats, ecosystems, and geopolitical boundaries (Barnosky et al., 2011; Wake & Vredenburg, 2008). Current extinction rates exceed those of the last five mass extinction events, with the sixth mass extinction event being driven by ever-growing anthropogenic activities (Ehlers & Krafft, 2006). The Anthropocene has been particularly detrimental to pollinators (Cameron et al., 2011; Hallmann et al., 2017; Lever et al., 2014; Macgregor et al., 2019; Potts et al., 2010), leading to numerous pollinator species being listed or petitioned for listing in the United States Federal Register (81 FR 67786, 82 FR 10285) as threatened or endangered. Pollinators are of vital importance to North American ecosystems and economies (Losey & Vaughan, 2006; Southwick & Southwick, 1992). The loss of pollinator biodiversity is of substantial concern as it reduces plant reproductive success (Klein et al., 2007; Ollerton et al., 2011; Thomann et al., 2013), erodes ecosystem services provided by pollination (Ollerton et al., 2011; Vanbergen & the Insect Pollinators Initiative, 2013; Winfree et al., 2011), and ultimately drives staggering economic losses (Gallai et al., 2009; Kevan & Phillips, 2001; Klein et al., 2007). Therefore, rapid, efficient, and accurate assessment of pollinator communities is a conservation imperative to inform adaptive management strategies to mitigate or prevent further loss of pollinator biodiversity (Carvalheiro et al., 2013; Didham et al., 2020; Montgomery et al., 2020; Thomas et al., 2019).

Such assessments are both technically difficult and financially costly (Bartomeus, 2013; Didham et al., 2020; Plein et al., 2017). Conventional monitoring is often restricted in terms of comparing multiple communities, requires high time investment to document presence or absence of pollinator species, and is reliant on dwindling taxonomic expertise to identify observed pollinators (Sheffield et al., 2009; Tur et al., 2013; Weiner et al., 2014). Moreover, monitoring the overwhelming majority of pollinator species is unrealistic (Didham et al., 2020; Young et al., 2017). These challenges are likely exacerbated for threatened and endangered species, whose populations tend to be numerically small and patchily distributed (Matthies et al., 2004). Yet, pollinator assessment is critical to monitor status and trends of rare, threatened and endangered pollinators, and evaluate the impacts of biodiversity loss on ecosystem services (Carvalheiro et al., 2013; Montgomery et al., 2020). Approaches that can estimate pollinator biodiversity in both a time-efficient and cost-effective manner will drastically enhance the development of effective interventions to mitigate or prevent further losses (Didham et al., 2020).

Environmental DNA (eDNA) analysis has emerged as a viable candidate for rapid, non-invasive biodiversity assessment (Deiner et al., 2017; Taberlet et al., 2012). Organisms deposit genetic material into their environment (e.g., via secretions and excretions) that can be harnessed to identify single species or entire communities in freshwater (Biggs et al., 2015; Broadhurst et al., 2021), marine (K. J. Harper et al., 2020; Valsecchi et al., 2021) and terrestrial (Clare et al., 2021; Katz et al., 2020) contexts. Single-species eDNA assays have been increasingly leveraged as a reliable means of surveying rare or invasive species (Thalinger et al., 2021). However, the costs associated with single-species assay development, validation and application, coupled with increasing numbers of imperiled species requiring monitoring (especially pollinators), necessitates methods that can simultaneously screen for multiple taxa, increasing efficiencies and reducing financial burdens (L. R. Harper et al., 2018; Moss et al., 2022; Wilcox et al., 2020). eDNA metabarcoding may provide a solution for more comprehensive and less costly assessment of invertebrate communities across larger spatial and temporal scales (L. R. Harper et al., 2020; Mächler et al., 2019; Roger et al., 2021).

eDNA metabarcoding is based on PCR amplification of short DNA fragments using primers with conserved binding sites across multiple taxa, flanking a region of highly variable DNA sequence that enables taxa to be distinguished from each other, followed by high-throughput sequencing (Deiner et al., 2017; Taberlet et al., 2012). The mitochondrial cytochrome c oxidase subunit I (COI) locus is the standard genetic marker for DNA barcoding of metazoans due to high variability among species (enabling high taxonomic resolution) and extensive reference database representation (Elbrecht et al., 2018). However, this variability has proven problematic for metabarcoding due to a paucity of conserved primer binding sites (Clarke et al., 2017; Leese et al., 2020). Degenerate primers can counter this issue but have inherent risks of primer mismatch, primer slippage, non-target amplification (e.g., bacteria, algae and fungi), and binding of non-target regions. All of these issues can introduce bias, complicating bioinformatic processing and taxonomic assignment (Clarke et al., 2017; Elbrecht et al., 2016, 2018; Leese et al., 2020).

Using different markers (e.g., 16S ribosomal RNA) or COI in conjunction with other markers has been investigated, but reference databases for other markers are comparatively sparse, limiting taxonomic resolution, and use of multiple markers increases PCR and sequencing costs (Clarke et al., 2017; Elbrecht et al., 2016; Elbrecht et al., 2017). Furthermore, good *in silico* performance of individual primer sets may not translate to *in situ* success, with performance also varying depending on the desired level of taxonomic resolution (Corse et al., 2019). Using multiple primer sets targeting the same marker has been proposed to combat this discrepancy in performance (Corse et al., 2019; Polanco F. et al., 2021), but does not avoid increased PCR and sequencing costs. Microfluidic technology may offer a solution by enabling PCR amplification with multiple primer sets targeting different taxa, different loci, or different regions of the same locus simultaneously, with all amplicons then being sequenced in single run (Brown et al., 2016). Hauck et al. (2019) demonstrated the potential of microfluidic eDNA metabarcoding to elucidate broader communities, provide high taxonomic resolution, and scrutinize performance of individual primer sets.

To our knowledge, only one study has evaluated eDNA metabarcoding of flowers as an approach to document pollinator communities (Thomsen & Sigsgaard, 2019). The authors successfully detected a wide range of terrestrial arthropods, including at least 135 species, but also highlighted issues of primer choice (despite pairing a COI primer set with a 16S primer set) and reference database representation. Our study builds upon this work using a combinatorial experimental design under both controlled laboratory and field conditions that tests the influence of flower species, sample type, and eDNA isolation protocol on pollinator communities detected by microfluidic eDNA metabarcoding. Based on our results, we propose a workflow that should enhance eDNA detection of pollinator communities and facilitate further research in this field.

## 2. Materials and methods

### 2.1 Greenhouse experiment

A mixed-species flowering plant assemblage was established in the University of Illinois Urbana-Champaign (UIUC) Plant Care Facility (Table S1; Figure 1a), hereafter greenhouse. Plantings were staggered so that a constant supply of pollen and nectar were available. A subset of flowering plants was moved to a secure room where a Natupol colony (Koppert Biological Systems) of common eastern bumblebees (*Bombus impatiens*) was released and allowed to acclimate for 10 days (Figure 1b). Temperature was set at 21-24°C during the day and 12-16°C during the night, humidity was ambient, and light was set to 14-hr day length.

**Figure 1.**
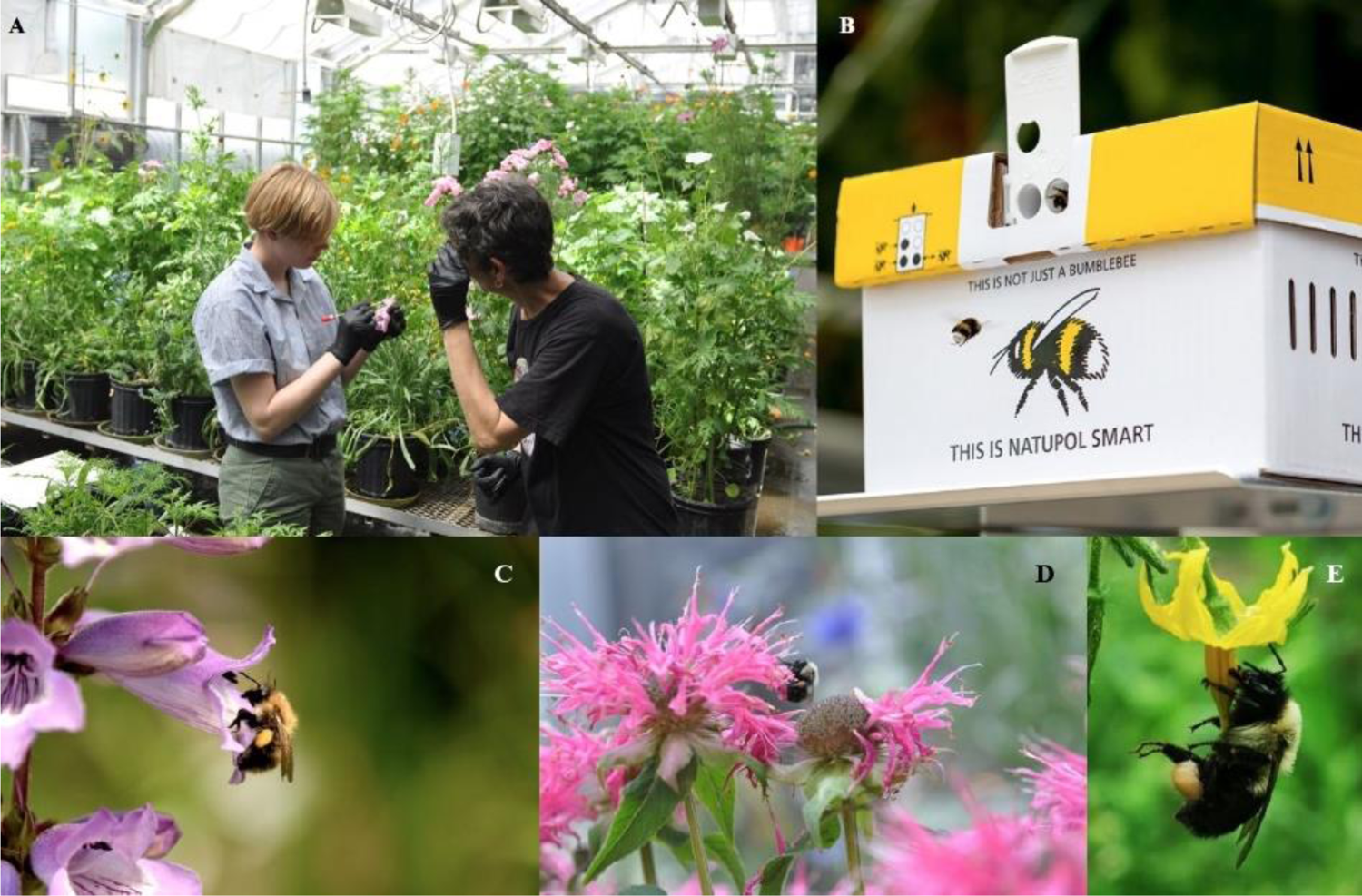
Images depicting greenhouse experiment and flower morphology, size, and characteristics of our focal flower species: *Penstemon* sp., *Monarda* sp., and *Solanum* sp.. We established a mixed-species colony of flowering plants to ensure a constant supply of pollen and nectar **(a)**. A subset of plants was then moved to an isolated greenhouse room where a colony of common eastern bumblebees was established **(b)**. The large, conical flowers of *Penstemon* **(c)** require pollinators to crawl in to access nectar/pollen. *Monarda* **(d)** inflorescences are composed of crowded, head-like clusters of flowers providing nectar and pollen. *Solanum* **(e)** inflorescences are considerably smaller and star-shaped with exposed stamens but no nectar. Photos by Mark Davis and Tiffany Jolley.

We selected four focal flower species with different flower morphologies to separately introduce to the secure room over two months: *Penstemon* sp., *Monarda* sp., *Solanum* sp., and *Cynoglossum amabile* (Figure 1c-e). Prior to the introduction of a flower species, flowers were sampled using up to three methods to provide a detection baseline for other insect species present in the greenhouse (baseline controls): 1) whole flower heads were harvested (*Penstemon*, *Monarda*, and *Solanum*), 2) the flower was swabbed with a flocked cotton swab (*Penstemon* and *Monarda* only), and 3) nectar was extracted from the flower via microcapillary tube (*Cynoglossum amabile* only). All three sampling approaches could not be applied to all four flower species because *Solanum* does not produce nectar and attempted nectar draws from *Penstemon* and *Monarda* were unsuccessful. Samples were preserved in Qiagen^®^ ATL buffer or cetyl trimethyl ammonium bromide (CTAB; Teknova) in sterile 2 mL microcentrifuge tubes (MidWest Scientific). Volumes of 300, 600, 900 or 1200 μL were used depending on sample size/type (see Table S2).

Upon introducing a flower species, we observed the flowers for bumblebee visits. After a visit by a bumblebee and any handling was observed, the flower was swabbed (*Penstemon* and *Monarda* only). A total of 50 flowers (one flower per plant) were sampled in one day, with 25 each preserved in ATL buffer and CTAB in sterile 2 mL microcentrifuge tubes. Flowers were left unobserved in the greenhouse for 24 hrs, then an approximately equal number of flowers from each of three benches were swabbed unsystematically to assess the ability to detect pollinator visits in the absence of direct observation. Again, 50 flowers (one flower per plant) were sampled in one day and divided equally between the two preservation buffers. This two-day process was repeated for harvesting of whole flower heads (*Penstemon*, *Monarda*, and *Solanum*), then nectar extraction from flowers via microcapillary tube (*Cynoglossum amabile* only). On each day that a flower species was sampled, sterile 2 mL microcentrifuge tubes containing 1200 μL of ATL buffer and CTAB were taken into the secure room but no sample was added. These tubes were transported, stored, and processed alongside samples (field blanks).

### 2.2 Field experiment

Two of the four focal flower species (*Penstemon* and *Monarda*) in the greenhouse also occurred on the UIUC campus enabling a pilot field component to the greenhouse experiment using a similar sampling approach. We concentrated field sampling effort on *Penstemon* first, followed by *Monarda* due to differences in timing of peak flowering. Flowers were observed between 9am and 12pm on days when the weather was suitable (sunny with temperatures >13°C). When a bee species visited a flower, the bee was identified (https://beespotter.org/) and a photo was taken. After the bee left the flower, the flower was swabbed. A total of 50 flowers (one flower per plant) were sampled in one day, with 25 each preserved in ATL buffer and CTAB in sterile 2 mL microcentrifuge tubes. On the next suitable day, 50 unsystematic swabs (i.e., no observation of pollinator visits) were acquired (one flower per plant) and divided equally between the two preservation buffers. This process was then repeated for harvesting of whole flower heads. Field blanks were included on each day that sampling occurred.

### 2.3 DNA extraction, quantification and plating

DNA extraction was performed at the Collaborative Ecological Genetics Laboratory, Illinois Natural History Survey, UIUC, where dedicated labs and established procedures for working with environmental samples of low DNA concentration are in place. Procedures include regular bleaching (50% v/v solution) of work surfaces, ultraviolet (UV) irradiation of reagents, consumables, and workspaces, isolated PCR hoods, and physical separation of pre-PCR (i.e., a dedicated eDNA “clean room” where no high-copy DNA has been introduced) and post-PCR processes.

A modified Qiagen^®^ DNeasy^®^ Blood & Tissue (QBT) protocol (Thomsen & Sigsgaard, 2019) was used for extraction of samples preserved in ATL buffer (see Supporting Information). Lysis was conducted in the tubes containing samples by adding 60, 100, 200, or 300 μL of proteinase K respectively depending on the volume of preservative used (Table S2). DNA was eluted in 2 × 60 μL of AE buffer, with a 15 min incubation step at 37°C each time before centrifugation. A modified Phenol-Chloroform-Isoamyl (PCI) extraction and ethanol precipitation method (Renshaw et al., 2015) was employed for samples preserved in CTAB (see Supporting Information). DNA was rehydrated with 100 μL of 1x TE buffer (ThermoFisher Scientific). Buffer-only extraction blanks were included for each batch of extractions (*n* = 23). DNA extracts were stored at −20°C until quantification and plating.

Samples extracted with both protocols were quantified using a Qubit™ 3.0 Fluorometer (Life Technologies, Grand Island, NY) with the Qubit™ dsDNA HS Assay Kit (ThermoFisher Scientific). Three readings were taken for each sample with the average used as the final concentration (ng/μL). We aliquoted 30 μL of each sample into a 96-well 0.2 mL optical reaction plate (ThermoFisher Scientific). A total of 42 baseline controls/eDNA samples, two field blanks, two extraction blanks, and two PCR positive controls (one of ten bee species known to occur in Illinois, listed in Figure 2 excluding *B. griseocollis*) were included on each plate. Plates were stored at −20°C until submission (typically less than 24 hrs) for microfluidic eDNA metabarcoding (see section 2.5).

**Figure 2.**
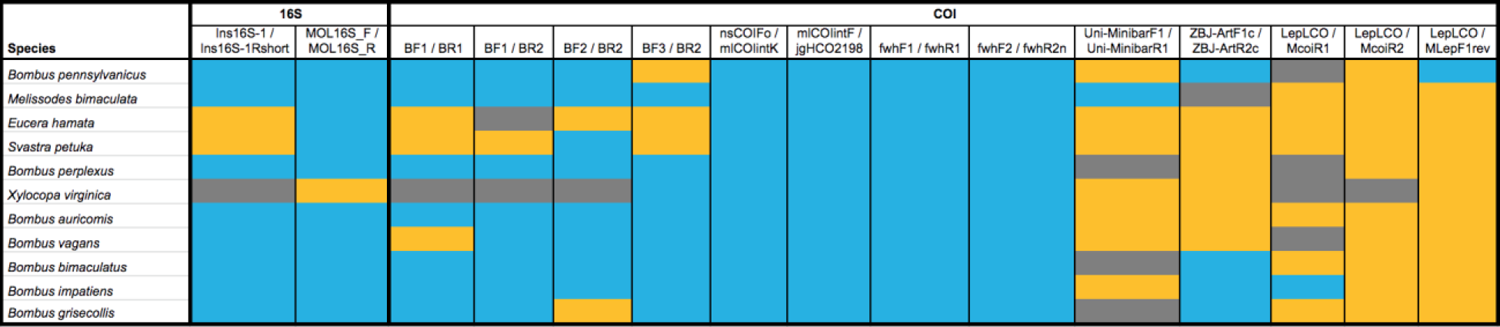
Tile plot depicting the results of *in vitro* primer validation. The subset of primers with approximately appropriate annealing temperature and fragment length were tested with 11 locally occurring bee species using PCR and verified via gel electrophoresis. Blue cells represent comparisons in which all three PCR replicates were positive, yellow cells represent comparisons in which one or two PCR replicates were positive, and grey cells represent comparisons in which all three PCR replicates failed.

### 2.4 Primer selection and validation

We conducted a comprehensive literature review to identify suitable primer sets for arthropod eDNA metabarcoding. We refined this list of primers to those with Fluidigm compatible annealing temperatures (∼55°C) that amplified a 150-400 bp fragment (lower bound to meet post-amplification removal of primer-adapted dimers, upper bound to limit amplicon length for paired-end sequencing) for microfluidic eDNA metabarcoding.

For *in silico* primer validation, we constructed a reference sequence database for invertebrate species that occur in Illinois using a list of binomial names based on observations by greenhouse staff, light-trapping data from the Illinois Bat Conservation Program (IBCP), arthropods and molluscs catalogued in the Illinois Natural History Survey (INHS) Insect Collection Database (http://inhsinsectcollection.speciesfile.org/InsectCollection.aspx), cicadas recorded by Catherine Dana (INHS), and threatened and endangered priorities provided by Angella Moorehouse (Illinois Nature Preserves Commission). With this list, all available sequences for markers amplified by the selected primer sets were downloaded from GenBank using Reprophylo v1.3 (Szitenberg et al., 2015) in July 2019. Reference sequences in GenBank format were converted to ecoPCR (Ficetola et al., 2010) database format using obiconvert (Boyer et al., 2016). ecoPCR parameters were set to allow a 50–500 bp fragment and up to 3 mismatches between each primer and each sequence in the reference database.

For *in vitro* primer validation, DNA extracts from 11 bee species known to occur in Illinois were amplified with the selected primer sets. (Figure 2). PCR was performed in triplicate using reaction volumes of 25 μL, including 12.5 μL of sterile molecular grade water (MidwestScientific), 11.1 μL of GoTaq^®^ Colorless Master Mix (ThermoFisher Scientific), 0.2 μL of forward primer and 0.2 μL of reverse primer (10 μM, Integrated DNA Technologies), and 1 μL of template DNA. Thermocycling conditions were 5 min at 94°C, followed by 40 cycles of 45 sec at 94°C, 1 min at 55°C and 1 min 30 sec at 72°C, then 5 min at 72°C. PCR amplification was confirmed using agarose gel electrophoresis (2 g agarose (Midwest Scientific), 100 mL 1x TAE buffer (Midwest Scientific)) at 120 V for 90 min, and gels were visualized with a LabNet Systems EnduroGel Gel Documentation System. Amplification was classed as successful (3 of 3 PCR replicates), moderately successful (1 or 2 of 3 PCR replicates), or unsuccessful (0 of 3 PCR replicates).

### 2.5 Microfluidic eDNA metabarcoding

Microfluidic eDNA metabarcoding, including PCR amplification, Illumina sequencing, and demultiplexing using QIIME 2™ (Bolyen et al., 2019), was conducted by the Roy J. Carver Biotechnology Center Functional Genomic Unit at the UIUC. The Fluidigm 48.48 Access Array™ (Fluidigm Corporation, 2016) was used for PCR amplification of each eDNA sample (2 ng) once with our validated primer panel (see Supporting Information). Products were quantified on a Qubit™ Fluorometer and stored at −20°C. All samples were run on a Fragment Analyzer (Advanced Analytics) and amplicon regions and expected sizes confirmed. Samples were then pooled in equal amounts according to product concentration. The pooled products were size selected on a 2% agarose E-gel (Life Technologies) and extracted from the isolated gel slice with a QIAquick Gel Extraction Kit (Qiagen). Purified products were run on an Agilent Bioanalyzer to confirm appropriate amplicon profile and determination of average size. The products were quantified using a Qubit™ Fluorometer, pooled evenly, and the final libraries were quantified by qPCR on a BioRad CFX Connect Real-Time System (Bio-Rad Laboratories, Inc.). The libraries were denatured, spiked with 20% PhiX Control v3, and loaded at 8 pM to an Illumina MiSeq for cluster formation and sequencing with a MiSeq Reagent Kit v3 (600-cycle) (Illumina, Inc.). In total, 40 baseline controls, 336 eDNA samples, 18 field blanks, 18 extraction blanks, two PCR negative controls, and 18 PCR positive controls were sequenced across two runs (288 samples/controls and 144 samples/controls, respectively). Sequences were sorted and demultiplexed by primer set and sample, then quality and yield assessed using the FastX Tool Kit (http://hannonlab.cshl.edu/fastx_toolkit/index.html).

### 2.6 Bioinformatic processing

The demultiplexed FASTQ files for each primer set were processed using metaBEAT v0.97.11 (https://github.com/HullUnibioinformatics/metaBEAT) and the Anacapa Toolkit v1 (Curd et al., 2019), which has been deposited on GitHub (https://github.com/limey-bean/Anacapa) and permanently archived (http://doi.org/10.5281/zenodo.3064152). Each pipeline incorporates a different suite of open-source software for bioinformatic processing (see Supporting Information), which allowed us to assess the robustness of eDNA detections. metaBEAT produced comparable or better detection than Anacapa for our focal pollinator species and species known to occur in the greenhouse (see Supporting Information and Figure S1), thus we used these results for downstream analyses.

### 2.7 Data analysis

All data manipulation and downstream analyses were conducted in R v3.6.3 (R Core Team, 2020). Datasets for each primer set were combined into a single master dataset, retaining primer information, and detections in eDNA samples and process controls were assessed (Figures S2, S3). Taxon richness within samples produced by each primer pair and taxon overlap between different primer pairs was assessed (Figures S4, S5) before removing primer information and merging reads for each taxon. A highly conservative false positive sequence threshold was calculated based on the maximum frequency of arthropod DNA (2.05%) found in the negative process controls (i.e., field blanks, extraction blanks, and PCR negative controls) and applied to the metabarcoding data. PCR positive controls were not used for threshold determination as DNA was isolated from bumblebee specimens sourced from pinned museum collections resulting in high contamination frequency, and the species that specimens belonged to could naturally occur in our study system (see Discussion). The data were filtered to remove non-arthropod taxonomic assignments before taxonomic assignments were filtered on a sample-by-sample basis to retain the lowest taxonomic ranks where assignments to multiple taxonomic ranks were present (e.g., species, genus and family). The filtered data were then combined with greenhouse and field metadata.

We examined taxa detected in the baseline controls before flower species were introduced to the secure room containing common eastern bumblebees (Figure S6), in eDNA samples after flower species were introduced to the secure room (Figure S7), and in eDNA samples taken from flowers around the UIUC campus (Figure S8). We then investigated biological and technical grouping variables that potentially influence pollinator eDNA detection: flower species, sample type, and eDNA isolation protocol. The read count data were converted to binary data as potential bias introduced by PCR amplification may prevent reliable abundance or biomass estimation from sequence reads produced by DNA or eDNA metabarcoding (Elbrecht et al., 2017).

First, alpha diversity (i.e., taxon richness) of eDNA samples according to each grouping variable was compared. The data were non-normally distributed and the assumptions of a one-way Analysis of Variance (ANOVA) were violated (Table S5, Figures S9, S10, S11), which Generalised Linear Models (GLMs) with different error families and link-functions did not resolve. Therefore, we compared taxon richness according to each grouping variable using the non-parametric Kruskal-Wallis test from the package stats v3.6.3, and performed multiple pairwise comparisons using the non-parametric Dunn’s test (p-values adjusted with the Benjamini-Hochberg method where applicable) from the package FSA v0.9.1 (Ogle et al., 2020).

The package betapart v1.5.4 (Baselga & Orme, 2012) was used to estimate total beta diversity, partitioned by nestedness-resultant (i.e., community dissimilarity due to taxon subsets) and turnover (i.e., community dissimilarity due to taxon replacement), across all samples with the *beta.multi*function. Greenhouse samples 80PEWF7-P2-G3, 80MOSW09-P3-C3, 80PEWF23-P1-G4 and 80PEWF73-P9-C5 and UIUC campus samples 80PESW105-P5-G2 and 80PEWF37-P2-A3 skewed visualisation of beta diversity components, thus these samples were removed from the dataset prior to performing these analyses. The three components of beta diversity (Jaccard dissimilarity) were estimated for eDNA samples according to each grouping variable using the *beta.pair* function. For each component of beta diversity, we compared the variance in each group of samples by calculating homogeneity of multivariate dispersions (MVDISP) using the *betadisper* function in the package vegan v2.5-7 (Oksanen et al., 2019). Differences in MVDISP between groups of eDNA samples were statistically tested using an ANOVA. Community dissimilarity for each component of beta diversity was visualized using Non-metric Multidimensional Scaling (NMDS) with the *metaMDS* function and tested statistically using permutational multivariate analysis of variance (PERMANOVA) with the function *adonis* in the package vegan. Pre-defined cut-off values were used for effect size, where PERMANOVA results were interpreted as moderate and strong effects if R^2^ > 0.09 and R^2^ > 0.25 respectively. These values are broadly equivalent to correlation coefficients of *r* = 0.3 and 0.5 which represent moderate and strong effects accordingly (Nakagawa & Cuthill, 2007). All plots were produced using the packages ggplot2 v3.3.5 (Wickham, 2016) and ggpubr v0.4.0 (Kassambara, 2020).

## 3. Results

### 3.1 Primer selection and validation

Our literature review revealed almost 120 individual primers that yielded over 60 primer combinations across two mitochondrial and two nuclear genes that were suitable for invertebrate metabarcoding, of which 22 were tested *in silico* (Table S3). A total of 15 primer sets designed to amplify regions of the mitochondrial COI and 16S genes were deemed suitable for microfluidic eDNA metabarcoding based on Fluidigm 48.48 Access Array™ parameters and taxonomic coverage (Table S3). These primer sets have been used in studies investigating diversity of terrestrial arthropods (Clarke et al., 2014), freshwater macroinvertebrates (Elbrecht & Leese, 2017; Vamos et al., 2017), freshwater molluscs (Klymus et al., 2017), and marine metazoans (Günther et al., 2018; Leray et al., 2013; Meusnier et al., 2008) as well as diet of various predators (Corse et al., 2019; Zeale et al., 2011). Several of the selected primer sets used here were recently vetted by Elbrecht et al. (2019) for terrestrial arthropod biodiversity assessment using DNA metabarcoding, where eight published primer sets outperformed 28 others that were tested, including: BF3/BR2 (Elbrecht et al., 2019), BF1/BR2 (Elbrecht & Leese, 2017), fwhF2/fwhR2n (Vamos et al., 2017), and mlCOIintF/jgHCO2198 (Geller et al., 2013; Leray et al., 2013). Our *in vitro* testing confirmed that the 15 selected primer sets successfully amplified pollinators, specifically 11 bee species that occur in Illinois (Figure 2). Given these results, we chose to retain the full primer panel for microfluidic eDNA metabarcoding.

### 3.2 Microfluidic eDNA metabarcoding

Each sequencing run generated 20,771,510 and 19,116,230 sequence reads, respectively. On average, each primer set produced 1,384,767 and 1,274,415 sequence reads for samples/controls on each sequencing run (average of 4,808 and 8,850 sequence reads per sample/control with each primer set). UniMinibarF1/UniMinibarR2 produced the fewest sequence reads (6,824 and 4,556) whereas mlCOIintF/jgHCO2198 (3,844,444 and 3,516,470) and Ins16S-1F/Ins16S-1Rshort (4,743,478 and 5,848,630) produced the highest reads for COI and 16S, respectively. The majority of taxonomically assigned reads belonged to Arthropoda for most primer sets, excluding BF2/BR2 (70.79% and 80.44%), BF3/BR2 (66.19% and 81.39%) and MOL16S_F/MOL16S_R (26.52% and 25.63%). The primer sets that proportionately had the most reads assigned to Apidae were nsCOIFo/mlCOIintK (2.01% and 14.11%), fwhF2/fwhR2n (2.91% and 7.94%), BF1/BR2 (1.23% and 3.72%), LepLCO/MLepF1rev (2.60% and 2.15%), and mlCOIintF/jgHCO2198 (1.98% and 3.60%). Raw read counts, bioinformatics parameters, taxonomically assigned read counts, and unassigned read counts are summarised for each primer set in Table S4.

Microfluidic eDNA metabarcoding correctly identified seven out of 10 bee species used as PCR positive controls to species level, with three species identified to genus or family level. However, the PCR positive controls contained a number of contaminants (Figure S3) due to the manner in which bee specimens were collected, stored, and preserved by the INHS Insect Collection. In contrast, the negative process controls (field, extraction, and PCR) exhibited very little contamination (Figure S3). Before false positive threshold application and filtering of taxonomic assignments, we detected 358 taxa in the 40 baseline controls and 336 eDNA samples. A total of 279 taxa were either consistently detected below the sequence threshold (2.05%) calculated using our negative process controls or non-invertebrate and of coarse taxonomic resolution. The final dataset after threshold application contained 79 taxa for downstream analyses.

### 3.3 Greenhouse experiment

The common eastern bumblebee was not detected in any baseline controls collected from flower species before they were introduced to the secure room (Figure S6), but was successfully detected in eDNA samples collected from flower species after their introduction. *Monarda* whole flowers (*n* = 1) and swabs (*n* = 3) preserved in CTAB yielded positive detections for the target species. All sample types from *Penstemon*, *Solanum*, and *Cyngolossum amabile* failed to detect the target species (Figure S7). However, *Solanum* whole flowers preserved in ATL buffer (*n* = 1) produced weak eDNA signals for common eastern bumblebee, but these were removed by our false positive sequence threshold.

In addition to the target species, pest control and resident arthropods were detected in samples collected from the greenhouse. Specifically, the silverleaf whitefly (*Bemisia tabaci*), dark-winged fungus gnat (*Bradysia impatiens*), western flower thrip (*Frankliniella occidentalis*), insidious flower bug (*Orius* spp.), and aphids (*Aphis* spp.) were detected (Figures S6 and S7). However, a number of beneficial insect species as well as species known to occur in the greenhouse were not detected, including *Amblyseius californicus*, *Brasysia coprophilia*, camel crickets (*Ceuthophilus* spp.), *Encarsia formosa*, convergent lady beetle (*Hippodamia convergens*), bold jumping spider (*Phidippus audax*), shore fly (*Scatella stagnalis*), spider mites (Tetranychidae spp.), banded-wing whitefly (*Trialeurodes abutiloneus*), and greenhouse whitefly (*T. vaporariorum*). Of these species, only shore fly was detected in *Cynoglossum amabile* nectar preserved in ATL buffer (*n* = 1) and *Penstemon* whole flowers preserved in CTAB (*n* = 2) before false positive threshold application. Nonetheless, several species not previously known to occur in the greenhouse were detected, for example, cigarette beetles (*Lasioderma serricorne*).

### 3.4 Field experiment

We observed flower visitation from seven bee species in the field: common eastern bumblebee, black-and-gold bumblebee (*B. auricomus*), two-spotted bumblebee (*B. bimaculatus*), brown-belted bumblebee (*B. griseocollis*), American bumblebee (*B. pensylvanicus*), European honey bee (*Apis mellifera*), and eastern carpenter bee (*Xylocopa virginica*). Prior to false positive sequence threshold application, the brown-belted bumblebee and half-black bumblebee (*B. vagans*) were detected at low frequencies from *Penstemon* whole flowers preserved in CTAB, but not *Penstemon* swabs or any *Monarda* samples. None of the observed bee species were detected after false positive sequence threshold application (Figure S8). This included the eastern carpenter bee despite widespread observations and extensive evidence of nectar robbing by this species (Figure S12). Arthropods that were not directly observed but were recovered via microfluidic eDNA metabarcoding included the eastern calligrapher (*Toxomerus geminatus*), margined calligrapher (*T. marginatus*), cigarette beetle (*Melanophthalma inermis*), banded garden spider (*Argiope trifasciata*), sheet weavers (*Islandiana flaveola* and *Tennesseellum formica*), house fly (*Musca domestica*), egg parasitoid wasps (*Telenomus podisi*), Asian tiger mosquito (*Aedes albopictus*), and firebrat (*Thermobia domestica*). Most taxa were detected from *Penstemon* whole flowers or swabs (Figure S8).

### 3.5 Influence of biological and technical factors on eDNA detection

Overall, flower species did not influence alpha diversity of eDNA samples collected from the greenhouse (H = 3.070, *P* = 0.381; Figure 3ai) or UIUC campus (H = 0.009, *P* = 0.925; Figure 3bi). Taxon richness for *Penstemon* tended to be higher than other greenhouse flower species (*Cynoglossum amabile*: Z = −1.457, *P* = 0.871; *Monarda* Z = −0.166, *P* = 0.868; *Solanum*: Z = 1.116, *P* = 0.529). Taxon richness for *Penstemon* was comparable to *Monarda* around the UIUC campus (Z = 0.094, unadjusted *P* = 0.925), but maximum taxon richness for *Penstemon* was higher. Beta diversity among greenhouse (Figure 3aii-iv) and field (Figure 3bii-iv) eDNA samples was largely driven by turnover as opposed to nestedness-resultant (Table 1). MVDISP did not significantly differ between flower species for any beta diversity component except nestedness-resultant in the UIUC campus samples (Table 1). Flower species had a moderate positive effect on turnover (Figure 3aii) and total beta diversity (Figure 3aiv) of greenhouse arthropod communities, and a weak positive effect on turnover (Figure 3bii) and total beta diversity (Figure 3biv) of UIUC campus arthropod communities (Table 1).

**Figure 3.**
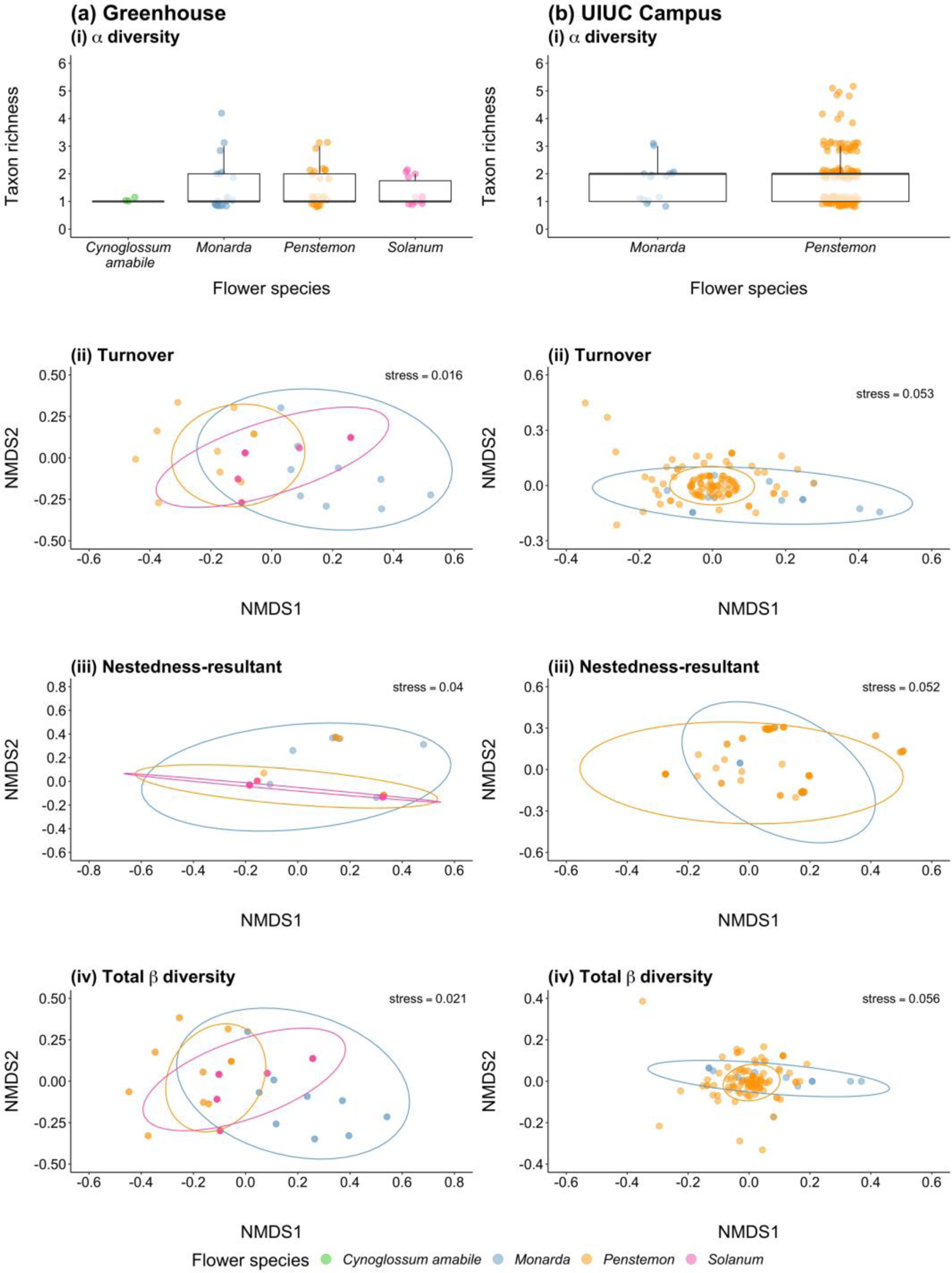
Summaries of alpha and beta diversity comparisons made between flower species sampled during **(a)** greenhouse and **(b)** field experiments: **(i)** boxplot showing the number of taxa detected in samples from each flower species, followed by non-metric multidimensional scaling (NMDS) plots of invertebrate communities for **(ii)** turnover, **(iii)** nestedness-resultant, and **(iv)** total beta diversity. Boxes show 25th, 50th, and 75th percentiles, and whiskers show 5th and 95th percentiles.

**Table 1.**
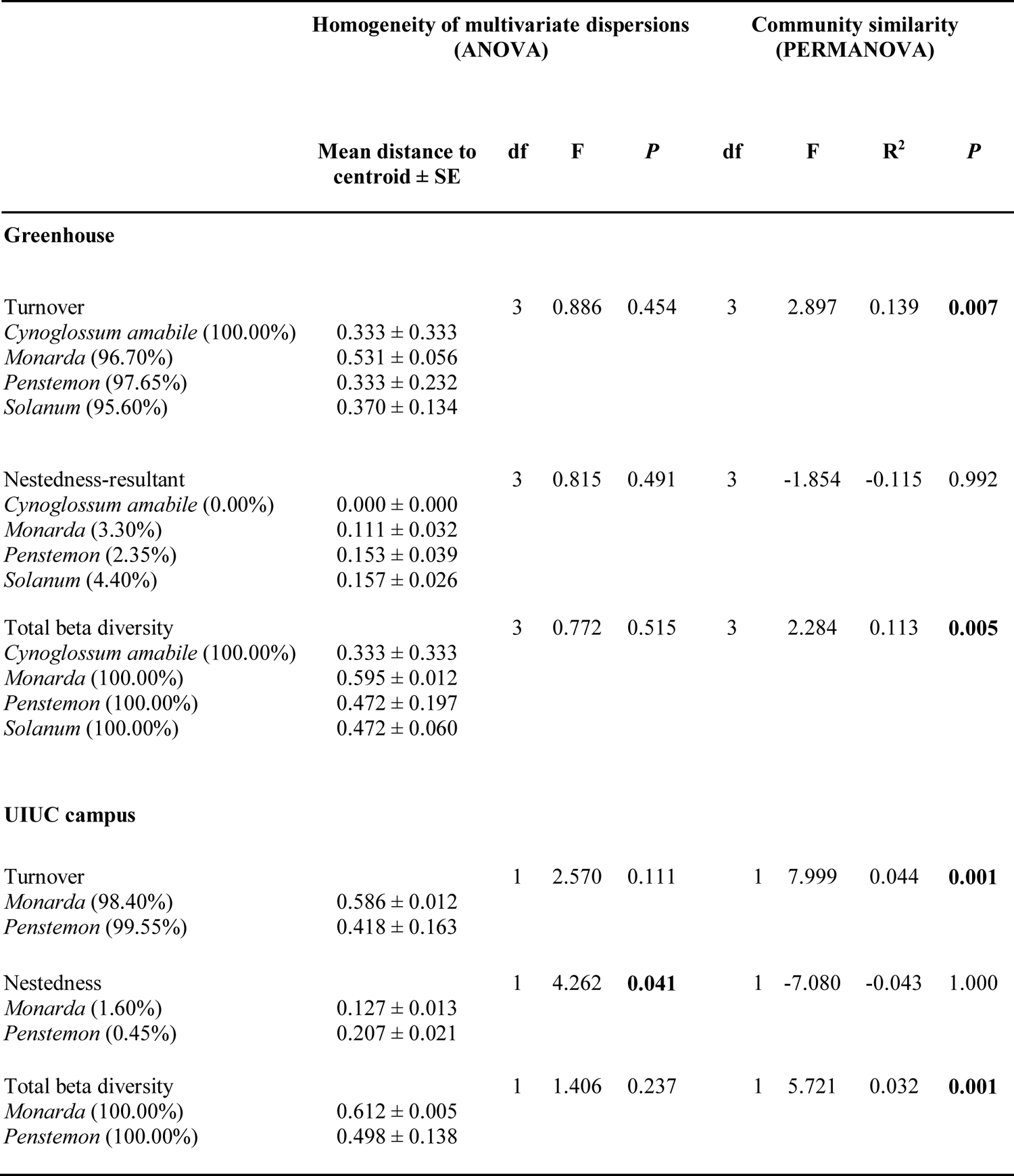
Summary of analyses statistically comparing homogeneity of multivariate dispersions (MVDISP) between the communities of different flower species (ANOVA) and variation in community composition of eDNA samples from different flower species (PERMANOVA). Relative contributions of taxon turnover and nestedness-resultant to total beta diversity (Jaccard dissimilarity) for each flower species are given in brackets. P-values in bold indicate statistical significance at 0.05.

Sample type did not influence alpha diversity of eDNA samples collected from the greenhouse (H = 3.923 *P* = 0.141; Figure 4ai), but did influence alpha diversity of UIUC campus eDNA samples (H = 6.459, *P* = 0.011; Figure 4bi). Taxon richness from nectar draws (*Cynoglossum amabile* only) was generally lower than swabs (Z = −0.692, *P* = 0.489) and whole flowers (Z = −1.494, *P* = 0.406) collected during the greenhouse experiment. In both the greenhouse and field experiments, swabs tended to produce lower taxon richness than whole flowers (greenhouse: Z = −1.473, *P* = 0.211; UIUC campus: Z = −2.541, unadjusted *P* = 0.011; Figures 4ai, bi). MVDISP did not differ between sample types for any beta diversity component except turnover in the UIUC campus samples (Table 2). Sample type had a weak positive effect on nestedness-resultant (Figure 4biii) of UIUC campus arthropod communities, but not any other component of beta diversity for greenhouse or UIUC campus samples (Table 2).

**Figure 4.**
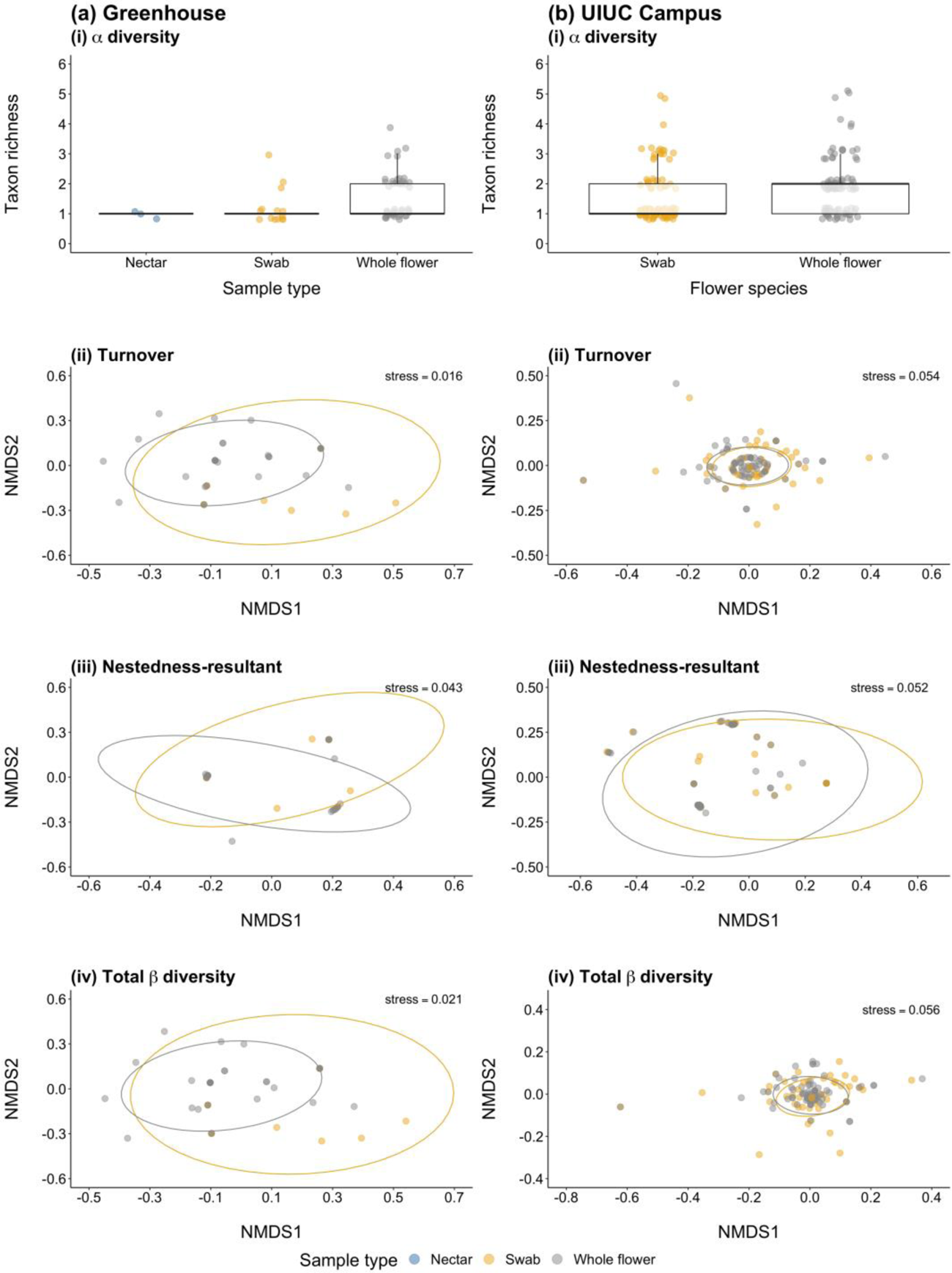
Summaries of alpha and beta diversity comparisons made between sample types collected during **(a)** greenhouse and **(b)** field experiments: **(i)** boxplot showing the number of taxa detected in samples from each sample type, followed by non-metric multidimensional scaling (NMDS) plots of invertebrate communities for **(ii)** turnover, **(iii)** nestedness-resultant, and **(iv)** total beta diversity. Boxes show 25th, 50th, and 75th percentiles, and whiskers show 5th and 95th percentiles.

**Table 2.** Summary of analyses statistically comparing homogeneity of multivariate dispersions (MVDISP) between the communities produced by different sample types (ANOVA) and variation in community composition of eDNA samples sourced from different plant material (PERMANOVA). Relative contributions of taxon turnover and nestedness-resultant to total beta diversity (Jaccard dissimilarity) for each sample type are given in brackets. P-values in bold indicate statistical significance at 0.05.

|  |  | Homogeneity of multivariate dispersions<br>(ANOVA) |  |  | Community similarity<br>(PERMANOVA) |  |  |  |
| --- | --- | --- | --- | --- | --- | --- | --- | --- |
| | Mean distance to<br>centroid ± SE | df | F | P | df | F | R<sup>2</sup> | P |
| **Greenhouse** |  |  |  |  |  |  |  |  |
| Turnover |  | 2 | 1.115 | 0.335 | 2 | 2.166 | 0.073 | 0.061 |
| Nectar (100.00%) | 0.333 ± 0.333 | | | | | | | |
| Swab (97.88%) | 0.582 ± 0.038 | | | | | | | |
| Whole flower (98.51%) | 0.397 ± 0.198 | | | | | | | |
| Nestedness |  | 2 | 1.816 | 0.172 | 2 | -0.402 | -0.015 | 0.816 |
| Nectar (0.00%) | 0.000 ± 0.000 | | | | | | | |
| Swab (2.12%) | 0.129 ± 0.012 | | | | | | | |
| Whole flower (1.49%) | 0.168 ± 0.029 | | | | | | | |
| Total beta diversity |  | 2 | 0.825 | 0.444 | 2 | 1.868 | 0.064 | 0.053 |
| Nectar (100.00%) | 0.333 ± 0.333 | | | | | | | |
| Swab (100.00%) | 0.591 ± 0.039 | | | | | | | |
| Whole flower (100.00%) | 0.508 ± 0.115 | | | | | | | |
| **UIUC campus** |  |  |  |  |  |  |  |  |
| Turnover |  | 1 | 4.730 | **0.031** | 1 | -0.950 | -0.006 | 0.999 |
| Swab (99.27%) | 0.333 ± 0.225 | | | | | | | |
| Whole flower (99.44%) | 0.469 ± 0.118 | | | | | | | |
| Nestedness |  | 1 | 0.307 | 0.580 | 1 | 13.989 | 0.075 | **0.002** |
| Swab (0.73%) | 0.160 ± 0.046 | | | | | | | |
| Whole flower (0.56%) | 0.176 ± 0.022 | | | | | | | |
| Total beta diversity |  | 1 | 1.478 | 0.226 | 1 | 0.774 | 0.005 | 0.654 |
| Swab (100.00%) | 0.478 ± 0.198 | | | | | | | |
| Whole flower (100.00%) | 0.544 ± 0.061 | | | | | | | |

eDNA isolation protocol did not influence alpha diversity of eDNA samples collected from the greenhouse (H = 0.012, *P* = 0.915; Figure 5ai) or the UIUC campus (H = 3.569, *P* = 0.059; Figure 5bi). There was very little difference in taxon richness between protocols for the greenhouse experiment (Z = 0.107, unadjusted *P* = 0.915; Figure 5ai). In the field experiment, taxon richness was typically greater in samples preserved in ATL buffer for QBT extraction (Z = 1.889, unadjusted *P* = 0.059) as opposed to CTAB preservation and PCI extraction (Figure 5bi). MVDISP did not differ between eDNA isolation protocols for any beta diversity component (Table 3). eDNA isolation protocol did not influence any beta diversity component (Table 3; Figures 5aii-iv, bii-iv).

**Figure 5.**
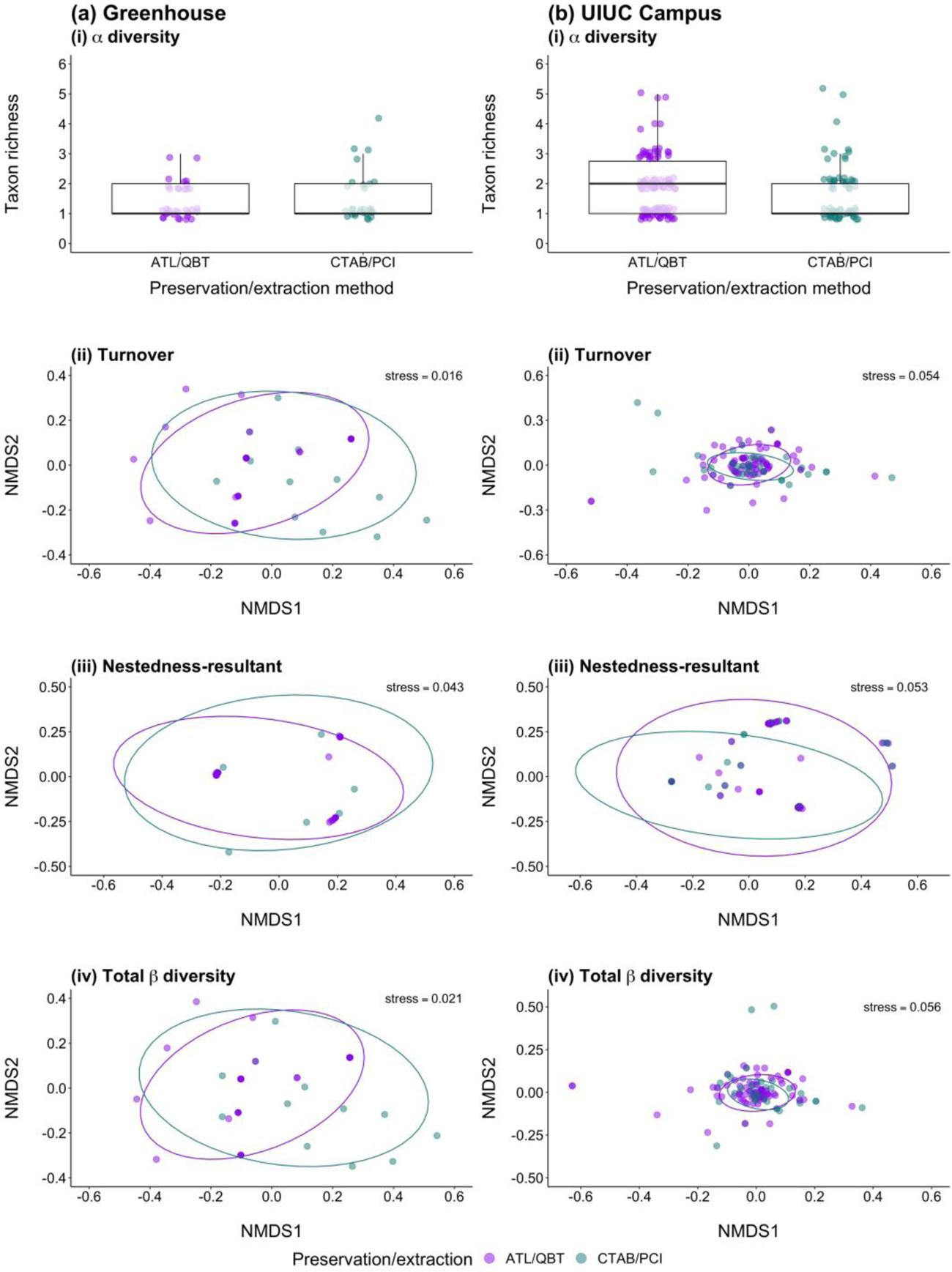
Summaries of alpha and beta diversity comparisons made between methods used for preservation and extraction of samples collected during **(a)** greenhouse and **(b)** field experiments: **(i)** boxplot showing the number of taxa detected in samples from each sample type, followed by non-metric multidimensional scaling (NMDS) plots of invertebrate communities for **(ii)** turnover, **(iii)** nestedness-resultant, and **(iv)** total beta diversity. Boxes show 25th, 50th, and 75th percentiles, and whiskers show 5th and 95th percentiles.

**Table 3.**
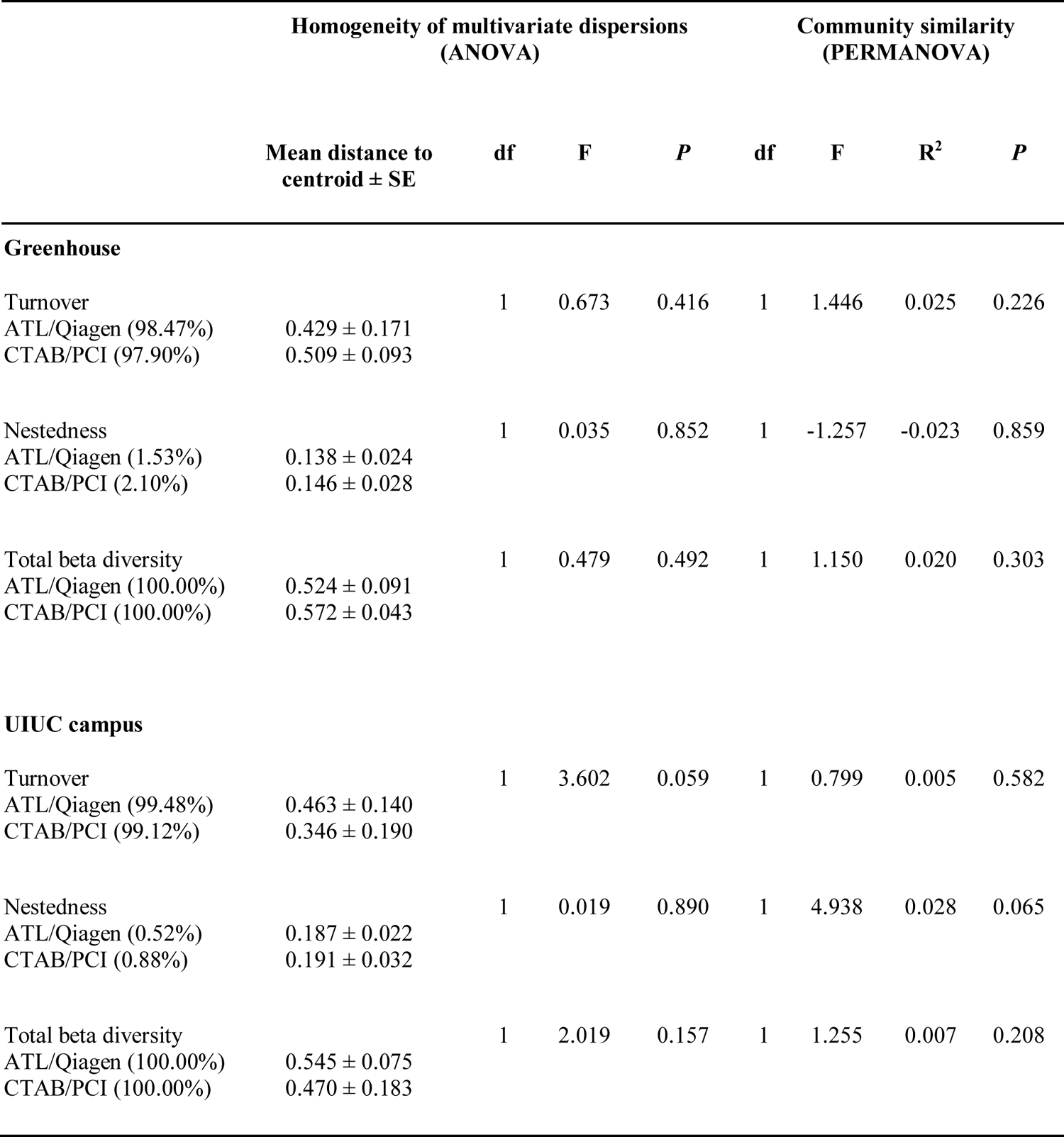
Summary of analyses statistically comparing homogeneity of multivariate dispersions (MVDISP) between the communities produced by different eDNA isolation protocols (ANOVA) and variation in community composition of eDNA samples preserved and extracted using different methods (PERMANOVA). Relative contributions of taxon turnover and nestedness-resultant to total beta diversity (Jaccard dissimilarity) for each eDNA isolation protocol are given in brackets. P-values in bold indicate statistical significance at 0.05.

## 4. Discussion

The past decade has yielded remarkable technological advances to address the critical need to execute rapid, large-scale surveys of pollinators using metabarcoding (Elbrecht et al., 2019; Roger et al., 2021; Thomsen & Sigsgaard, 2019). More recently, microfluidic eDNA metabarcoding has enabled researchers to “cast a broader net” and compensate for taxonomic biases or blind-spots inherent to individual primer sets (Hauck et al., 2019). We applied microfluidic eDNA metabarcoding to flower samples and found this approach to be a viable means of documenting arthropod communities, including pollinators, confirming the work of Thomsen & Sigsgaard (2019). Moreover, we tested biological and technical sources of variability in arthropod eDNA detection to develop an optimized workflow for eDNA-based pollinator biodiversity assessment. Below we contextualize our results and provide a framework for subsequent field research aimed at further validation of eDNA metabarcoding for assessment of all pollinator groups (e.g., insects, bats, and birds).

### 4.1 Greenhouse and field experiments

Both experiments provided important insights for field survey design. The most notable result of our greenhouse experiment was that microfluidic eDNA metabarcoding was effective in detecting the target common eastern bumblebee from one of our focal flower species (*Monarda*). We also detected insects that had been released for pest control and residents known to occur in the greenhouse facilities, further underscoring the utility of the approach to document arthropod communities. However, detection was imperfect as only *Monarda* swabs and whole flowers preserved in CTAB for PCR extraction yielded positive bumblebee detections, and several beneficial insect species as well as species known to occur in the greenhouse were not detected. We drew nectar from, swabbed, and harvested multiple flowers from each focal species immediately after visits from bumblebees, but *Penstemon*, *Solanum*, and *Cynoglossum amabile* did not yield any detections. The positive *Monarda* detections were flowers that had sat in the greenhouse for prolonged periods of time, suggesting that eDNA may accumulate to high enough levels for detection only after repeated visits by pollinator species, similar to other terrestrial species and aquatic or soil eDNA (Kucherenko et al., 2018; Williams et al., 2018).

Expanding upon the greenhouse experiment, eDNA metabarcoding detected pollinators and other plant-associated arthropods in natural flower plots around the UIUC campus, though again with interesting caveats. Numerous arthropod taxa were detected from our focal flower species (*Monarda* and *Penstemon*), including Arachnida, Coleoptera, Diptera, Hymenoptera, and Lepidoptera. Yet, not all observed bumblebee species were detected, suggesting that inherent characteristics of some pollinator species and/or their interactions with various plant morphologies may make them difficult to detect with this approach. Conventional surveys (e.g., visual, acoustic, and trapping) of arthropod communities that incorporate expert taxonomic identification would be needed for comprehensive estimation of non-detection rates and to pinpoint taxa which are frequently missed by eDNA metabarcoding.

Nevertheless, our controlled greenhouse experiment and pilot field experiment suggest that the development of an intensive sampling regime using flower swabbing and whole flower harvest across multiple flower species differing in their morphology, flowering phenology, etc. at fine spatiotemporal scales would yield a robust approach to rapidly assess pollinator communities in diverse ecosystems. We expand upon the measures needed to advance eDNA-based pollinator biodiversity assessment below.

### 4.2 Influence of biological and technical factors on eDNA detection

Thomsen and Sigsgaard (2019) provided the first evidence that eDNA metabarcoding could be used to monitor pollinator communities and other plant-associated arthropods by sequencing wild flowers. Our study advanced this monitoring approach by evaluating the influence of methodological choices throughout the eDNA metabarcoding workflow on the pollinator communities detected. Specifically, we tested four flower species with distinct morphological characteristics, three sample types, and two eDNA isolation protocols. While not extensively compared here, we also used 15 primer sets for two markers and processed our data using two bioinformatics pipelines. Our results show that eDNA detection of pollinator communities (and other plant-associated arthropods) is likely driven by synergistic or antagonistic effects of biological and technical factors. While differences in alpha diversity were negligible, community composition was influenced by flower species and sample type in both the greenhouse and field experiments. Taxa detected from one flower species were different taxa to those detected from other flower species, but taxa detected from one sample type were subsets of the communities detected with other sample types. These differences may arise from several sources, but suggest the diversity of arthropod communities revealed is likely dependent upon the diversity of the plant community sampled.

Flower morphology likely affects the ability of flower species to capture and retain eDNA from pollinators. Three of our focal flower species have distinct morphologies (Figures 1c-e). *Penstemon* has deep, conical flowers that require insects to crawl into the flower to seek pollen and nectar. *Monarda* has multiple small flowers arranged in a head-like cluster providing pollen and nectar. *Solanum* flowers are considerably smaller, with petals that do not obscure the stamen and no nectar. We hypothesize that the small size, low surface area, and exposed stamens of *Solanum* flowers coupled with the buzz pollination strategy required for this plant resulted in limited contact and thus less DNA deposition from the target common eastern bumblebee in the greenhouse, ultimately lowering detection probability. Although common eastern bumblebee DNA was detected in a *Solanum* sample, the signal was weak and subsequently removed by our false positive sequence threshold. In contrast, *Monarda* inflorescences are large, complex, have more overall nectar, and thus have a morphology that promotes longer handling and consequently increased deposition of DNA and probability of capturing and retaining pollinator eDNA. *Penstemon* has similar potential, as inflorescences are deep, ostensibly resulting in more pollinator surface area contact, and flower surfaces less exposed to the elements. Environmental exposure (e.g., temperature, rainfall, UV, and microbial activity) was previously found to lower detection of prey items from faeces with metabarcoding due to faster DNA degradation (McInnes et al., 2016; Oehm et al., 2011), and could similarly affect eDNA on flowers.

Morphological differences may also lead to differential handling times for insect species. Though we did not quantify handling time in this study, our observations indicate that bumblebees spent considerably longer on *Monarda* inflorescences, spent less time on *Penstemon* flowers, and spent the least amount of time on *Solanum* flowers. Our observations echo numerous studies that have identified differential handling times by bumblebees across a variety of flowers (Franco et al., 2011; Ivey, Martinez, & Wyatt, 2003), which is associated with morphological variation among bumblebee species (Harder, 1983). Increased contact and handling time with flowers may confer higher detection probabilities, ultimately influencing the ability of eDNA metabarcoding to recover pollinator communities. Curiously, some species that were frequently observed left no trace of eDNA on flowers, such as eastern carpenter bees. In *Penstemon* flower beds on the UIUC campus, we observed numerous visits by carpenter bees and collected multiple samples immediately after visitation. Moreover, nearly every flower we inspected showed evidence of nectar robbing (Figure S12), yet no reads were assigned to this species before or after false positive threshold application. Eastern carpenter bee DNA was used as a PCR positive control and successfully detected, thus lack of detection was not due to PCR, sequencing, or bioinformatics failure.

Sample type also influenced pollinator community composition. Sampling the entire flower head, irrespective of flower species, produced more species detections than swabbing or nectar draws. This method has already been established as an effective means of documenting pollinator communities (Thomsen & Sigsgaard, 2019), and likely results in improved chance of eDNA capture as well as increased DNA yield. Nevertheless, swabbing was able to detect pollinators, including the target common eastern bumblebee, and may be a more appropriate means of eDNA sampling in remote locations where it is unreasonable to transport the requisite sampling materials for whole flower harvesting, or in situations where a flower species is rare and/or legally protected. Nectar draws may be a less practical method to detect eDNA from pollinators for a number of reasons. First, some flowering plants do not produce nectar (e.g., *Solanum*), limiting the plant species that could be sampled with this approach. Second, nectar robbing may render nectar-producing flowers unable to be sampled due to nectar depletion. Finally, volumes of nectar among flower species can be variable and, in many instances, incredibly low, which may reduce detection probability. The effect of interplay between nectar depletion by a pollinator and nectar regeneration by a plant on eDNA detection is also unknown. Further exploration with high volume nectar-producing plants pollinated by insects as well as bird- and bat-pollinated plants is critical to close this knowledge gap.

eDNA isolation protocols are known to substantially influence eDNA detection of target taxa (e.g., Djurhuus et al., 2017; Hinlo et al., 2017; Renshaw et al., 2015; Spens et al., 2016), but comparisons have largely focused on aquatic eDNA and may not be transferable to eDNA on flowers (but see Lear et al., 2018). In our study, eDNA isolation protocol did not influence pollinator eDNA detection. Overall, we found ATL preservation with a modified QBT extraction (Thomsen & Sigsgaard, 2019) generally recovered greater taxon richness than CTAB preservation and PCI extraction (Renshaw et al., 2015), but the latter was more effective in detecting our target bumblebee species in the greenhouse experiment. Community dissimilarity stemmed from communities produced by CTAB preservation and PCI extraction being comprised of different taxa to the communities produced by ATL preservation and QBT extraction. These results reaffirm findings from aquatic eDNA research (Djurhuus et al., 2017; Hinlo et al., 2017; Lear et al., 2018), but further investigations using samples from natural flower plots where species inventories from conventional surveys are available would be worthwhile.

We used microfluidic eDNA metabarcoding to counteract some of the biases introduced in standard eDNA metabarcoding by primer choice. When performance of individual primer sets was examined, there was substantial variation in the total number of taxa within (Figure S4) and across (Figure S5) samples as well as the number of unique taxa detected by each primer set and distinct taxa shared by different combinations of primer sets (Figure S5). This reinforces the importance of carefully evaluating primer sets *in silico* and *in vitro*, and using primers targeting different markers and different regions (Corse et al., 2019; Polanco F. et al., 2021). Where studies are limited to selection of one primer set or marker for budgetary reasons, COI primers maintain their position in providing greater taxonomic coverage and resolution for arthropods, specifically mlCOIintF/jgHCO2198 (Geller et al., 2013; Leray et al., 2013), ZBJArtF1c-ZBJArtR2c (Zeale et al., 2011), BF3-BR2 and BF2-BR2 (Elbrecht et al., 2019; Elbrecht & Leese, 2017). However, metabarcoding is an evolving field with new primers targeting established regions as well as new markers targeting novel regions constantly emerging, and the best primers and markers will likely vary with target taxonomic group. Therefore, we advocate for continued assessment of individual primer sets and primer set combinations to maximize the biodiversity recovered whilst minimising effort and costs associated with eDNA metabarcoding.

Similarly, different bioinformatics pipelines can produce variable results due to the different softwares, parameterizations and algorithms implemented. We assessed two bioinformatics pipelines using different tools, namely clustering (metaBEAT) versus ASV parsing (Anacapa) and a custom reference database (metaBEAT) versus a CRUX database (Anacapa) (Curd et al., 2019; Hänfling et al., 2016). metaBEAT detected more arthropod species known to occur in the greenhouse facilities (Figure S1), thus data produced by this pipeline were used for downstream analyses. As bioinformatics pipelines continue to be developed and refined for processing metabarcoding data and public reference databases are bolstered by additional sequences, continued evaluation of bioinformatics pipelines is necessary to assess potential biases and taxon recovery (Mathon et al., 2021; O’Rourke et al., 2020). As such, we recommend that future studies assess different bioinformatic pipelines and their effects on species detection against known species inventories to help minimise the occurrence of false negatives and false positives in eDNA metabarcoding datasets.

### 4.3 Using hindsight to inform foresight

In our study, samples were PCR amplified and sequenced once. This was likely a main factor contributing to low eDNA detection of plant-associated arthropods present in the greenhouse facilities and observed in natural flower plots as there were no additional PCR replicates to offset PCR stochasticity and enable an additive strategy to bolster species inventories (Alberdi et al., 2018). Typically, metabarcoding studies have used three PCR replicates that are either pooled for sequencing or sequenced independently (Alberdi et al., 2018; Beentjes et al., 2019; Weigand & Macher, 2018), but eight or more PCR replicates may be required when detection probabilities are likely to be low (Ficetola et al., 2015). Eight PCR replicates were used by Hauck et al. (2019) for microfluidic eDNA metabarcoding, but some standard eDNA metabarcoding studies have used as many as 12 (Valentini et al., 2016) or 15 (Sato et al., 2017) PCR replicates. Greater PCR replication can improve detection rates for rare and/or low-density species, and enable identification of potential false positives if taxa occur in only one PCR replicate. However, removal of taxa occurring in a single replicate may increase the possibility of false negatives for rare and/or low-density taxa (Weigand & Macher, 2018). Conversely, some studies have suggested there may be little impact of including taxa that occur in only one PCR replicate on inferences of biodiversity patterns and community dissimilarly (Beentjes et al., 2019). Therefore, PCR replication may be of greater importance in alpha diversity estimation than beta diversity estimation. This is a key area of investigation for future pollinator and plant-associated arthropod eDNA research.

Similarly, more biological replication may be necessary to enhance pollinator eDNA detection rates, as has been observed for eDNA detection of terrestrial vertebrates from water (Broadhurst et al., 2021; T.-H. Macher et al., 2021). Here, only one flower was sampled per plant, but plant-associated arthropods may not have interacted with this particular flower. Since this study was conceived, alternative strategies for eDNA detection of terrestrial insects have been successfully trialled, such as eDNA aggregation via water (Valentin et al., 2020), airborne eDNA (Roger et al., 2022), and eDNA deposited on leaves (Krehenwinkel et al., 2022). These approaches may circumvent the need for biological replication and invasive sampling of flowers but require extensive validation against the current sampling strategies of swabbing flowers and/or whole flower harvest for pollinator eDNA used here and by Thomsen and Sigsgaard (2019).

Another important consideration is the use of microfluidic technology. High sequencing depth is key alongside PCR replication to improve detection of rare and/or low-density taxa in complex environmental samples (Alberdi et al., 2018; Doble et al., 2019; Hajibabaei et al., 2019). However, sequencing depth is inherently lower with microfluidic eDNA metabarcoding as it is split across the number of primer sets being used. Although we sequenced eDNA samples across two MiSeq runs, sequencing depth may have been insufficient for our primer panel consisting of 15 primer sets. There is an important trade-off to consider, as using a larger primer panel with the Fluidigm 48.48 Access Array™ allows the performance of more individual primer sets to be evaluated. Prior to false positive threshold application, we found that the primer set mlCOIintF/jgHCO2198 (Geller et al., 2013; Leray et al., 2013) detected the most taxa, followed by BF2/BR2 and BF3/BR2 (Elbrecht et al., 2019; Elbrecht & Leese, 2017), but each of the 15 primer sets detected taxa not identified by other primer sets, and different combinations of primer sets detected taxa not identified by other combinations (Figure S5). More sequencing depth is possible with the use of more powerful sequencers, such as the Illumina NovaSeq, with cost of these sequencing platforms being the main barrier to uptake at present. However, as the cost of Illumina sequencing continues to fall, greater sequencing depth can be achieved at equivalent or lower price points (even compared to one year ago). A magnitudinal increase in reads returned should also increase the likelihood of detecting DNA traces present at lower concentrations because they were left behind by species that are rarer, smaller, present at lower density or exhibited less flower handling time. This would greatly improve the efficacy of pollinator eDNA metabarcoding. However, where rare pollinators are the focus of monitoring efforts, it may be more appropriate to use targeted eDNA analysis (e.g. qPCR) for increased sensitivity (L. R. Harper et al., 2018; Moss et al., 2022).

PCR reaction volumes are also much lower using the Fluidigm 48.48 Access Array™ (5 μL) than they would be in standard PCR (typically 25 μL or 50 μL). Consequently, the amount of template DNA added to Fluidigm 48.48 Access Array™ reactions is a fraction of that used in standard PCR reactions, which is problematic as higher volumes of template DNA reduce uncertainty in detection rates (Mächler et al., 2015). In our study, 2 ng of template DNA was loaded to the Fluidigm 48.48 Access Array™, providing 0.13 ng DNA for amplification with each primer set in each reaction chamber. Although the initial volume of template DNA could be increased, the risk of inhibition or overwhelming PCR reactions also increases (Lance & Guan, 2020). Furthermore, the Fluidigm 48.48 Access Array™ must be run at a fixed annealing temperature which may be suboptimal for certain primer sets, thus impairing species detection (Clarke et al., 2017; Elbrecht et al., 2019). Therefore, metabarcoding projects may wish to exploit microfluidic technology in the first instance to determine which primer sets maximise community diversity, provide the most informative data, and should be used for processing of future samples (Brown et al., 2016), whilst acknowledging the caveats associated with this technology.

Contamination of eDNA samples can arise from many sources, including field sampling, DNA extraction, PCR, or sequencing errors (Ficetola et al., 2015; Ficetola et al., 2016; Sepulveda et al., 2020). We identified low levels of contamination in field and extraction blanks as well as PCR negative controls, yet considerably higher levels of contamination in PCR positive controls. The specimens used for our PCR positive controls were collected using pan traps, bulk preserved in ethanol, and sorted by hand without sterile gloves in unsterile conditions. The samples were then pinned and stored using standard entomological practices in open drawers at the INHS for several years. Given the above, we speculate that the frequency and composition of contaminants (including other pollinators, mites, and decomposers) likely occurred from contact with: 1) other species in the field prior to sampling (pre-catch), 2) other species during sampling and preservation (catch), and/or 3) common arthropod species living in museum collections (post-catch). This contamination led to the implementation of an ostensibly high and exclusionary false positive sequence threshold based on our negative process controls which likely removed true detections. Therefore, careful selection and procurement of specimens and their DNA for PCR positive controls will reduce threshold stringency, further enhancing the performance of pollinator eDNA metabarcoding.

## 5. Conclusions

Our study adds to a growing body of evidence that eDNA metabarcoding is a viable means of surveying pollinator communities. Methodological choices throughout the entire eDNA metabarcoding workflow, from what to sample, how to sample, eDNA isolation protocol, primer choice, and bioinformatics pipeline, can impact the community diversity elucidated using this approach. Based on our results, we have synthesised a recommended workflow for pollinator biodiversity assessment using eDNA metabarcoding (Figure 6). First, given the observed differences in detection across flower species, we recommend that replicate samples from a diverse flower species assemblage from a given site be established. By maximizing flower diversity (and considering spatial and temporal variation in pollinator communities), one should maximize the probability of detecting a comprehensive pollinator (and other plant-associated arthropod) community. Second, to maximize detection of diverse arthropod/pollinator communities, we recommend that whole flower heads are sampled. This increases the surface area available for eDNA to be captured from and may require less time to sample than swabbing or nectar draws. However we note that in certain contexts (i.e., in remote locations and/or in dealing with threatened/endangered plants that cannot be destructively sampled) swabbing may be a viable alternative. Third, ATL preservation followed by a modified QBT extraction generally revealed more diverse pollinator communities. Moreover, this approach tends to recover clean, long-stranded DNA. Fourth, the COI locus performed better than the 16S locus in recovering pollinator communities and minimizing the number of unassigned reads due to greater reference database coverage. Additionally, use of multiple primer sets helped minimize taxonomic biases of individual primer sets. On this basis, we recommend using a suite of COI primers in future metabarcoding studies of insect pollinators, but this is unlikely to be optimal for other pollinator groups (e.g., bats and birds). As such, *in silico* and *in vitro* primer validation should be undertaken for available primer sets for common metabarcoding markers (e.g., COI, 12S rRNA, 16S rRNA, 18S rRNA) depending on the pollinator community one seeks to document. Finally, metaBEAT seemingly recovered more diversity and detected more species known to be present in greenhouse facilities than Anacapa, but results may have been influenced by the use of different reference databases and tools for bioinformatic processing. Therefore, we advise future studies should undertake comparisons with available pipelines and compare data to known species inventories to select the most appropriate pipeline for their dataset.

**Figure 6.**
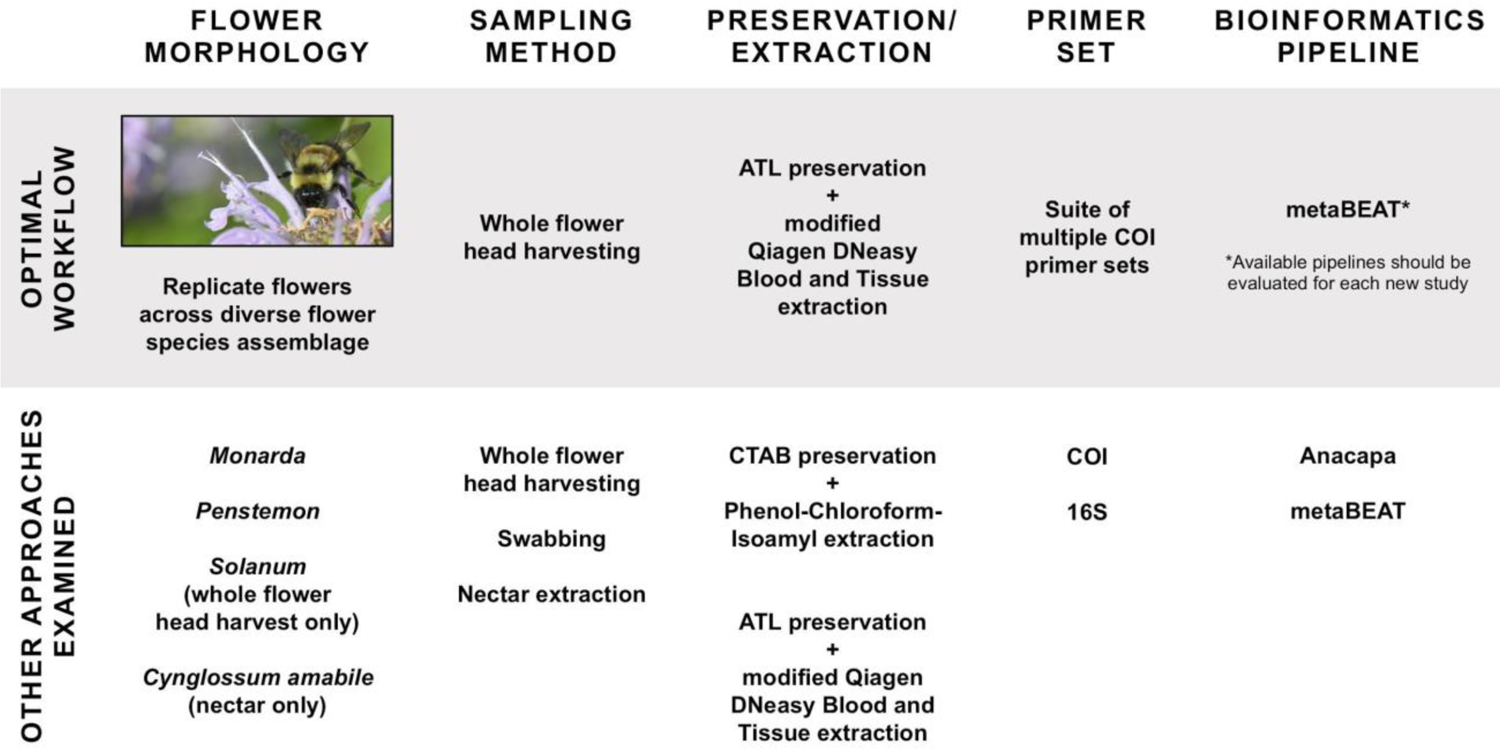
A recommended workflow for pollinator eDNA metabarcoding. This workflow was developed based on the results of this study and includes technical recommendations that should yield the most robust estimate of pollinator communities.

## Acknowledgements

We are grateful for the Strategic Environmental Research and Development Program for funding this research (grant number RC19-1102). We thank Patricia Dickerson for technical assistance. We are deeply grateful to Rosalie Metallo, Montgomery Flack, and Clinton Shipley at the UIUC Plant Care facility for technical, logistical, and moral support of our research. We appreciate Dr. Mark R. Band, Dr. Alvaro G. Hernandez, Dr. Christy L. Wright and the staff at the Roy J. Carver Biotechnology Center Functional Genomics Unit for technical support and conducting microfluidic metabarcoding. We thank Emily Curd for assistance with implementing Anacapa. We recognize Danielle Ruffatto, Tiffany Jolley, and Joe Spencer for imaging and design support. We thank Catherine Dana, Tara Hohoff, and Tommy McElrath for providing samples, as well as encouragement and creative criticism to improve this research. We thank Angella Moorehouse at the Illinois Nature Preserves Commission for supplying species lists used to develop the reference DNA sequence database.

## Authors contributions

M.A.D., B.M-F., and M.L.N. conceived and designed the study. B.M-F., L.E.P., and E.K. established the greenhouse flower assemblage and M.A.D., L.E.P., E.K., and B.M-F. collected samples. M.A.D., L.E.P., and E.K. performed laboratory work before samples were sent to the sequencing facility. M.L.N., J.B., and L.R.H. performed bioinformatics processing, and L.R.H. analysed the data. L.R.H. and M.A.D. wrote the manuscript, which all authors revised.

## Data availability statement

Raw sequence reads, bioinformatics scripts, R scripts, and corresponding data have been permanently archived at Zenodo (https://doi.org/10.5281/zenodo.6756972).

