## Supporting Information for "BeeDNA: microfluidic environmental DNA metabarcoding as a tool for connecting plant and pollinator communities"

#### Contents

**Methods** 3
DNA extraction 3
Microfluidic eDNA metabarcoding 4
Bioinformatic processing 4
**Tables** 7
Table S1 7
Table S2 8
Table S3 9
Table S4 12
Table S5 13
**Figures** 14
Figure S1 14
Figure S2 15
Figure S3 16
Figure S4 17
Figure S5 18
Figure S6 20
Figure S7 21
Figure S8 22
Figure S9 23
Figure S10 24
Figure S11 25
Figure S12 26
**References** 27

#### Methods

*DNA extraction*

A modified Qiagen® DNeasy® Blood & Tissue (QBT) protocol (Thomsen & Sigsgaard, 2019) was used for extraction of whole flower heads, swabs, and nectar draws preserved in ATL buffer. Lysis was conducted in the tubes containing samples by adding 60, 100, 200, or 300 µL of proteinase K respectively depending on the volume of preservative used (Table S2). Samples were agitated at 56°C in a rotor for 3 hrs. After lysis, samples were vortexed for 10 sec and 360, 700, 1,100, or 1,500 µL of lysate was retrieved, respectively. Equal amounts of AL buffer and absolute ethanol (EtOH; ThermoFisher Scientific) corresponding to the volume of retrieved lysis mixture were added to the tubes and vortexed thoroughly. The lysate from each sample (700 µL) was added to a spin column and passed through the membrane filter using centrifugation. Up to three centrifuge iterations were required to process all lysate. Spin columns were washed by adding 600 µL of AW1 buffer then 600 µL of AW2 buffer. Finally, DNA was eluted in 2 × 60 µL of AE buffer, with a 15 min incubation step at 37°C each time before spinning. All centrifugations were performed at 10,000 × g.

A modified Phenol-Chloroform-Isoamyl (PCI) extraction and ethanol precipitation method (Renshaw et al., 2015) was employed for whole flower heads, swabs, and nectar draws preserved in CTAB. CTAB (450 µL) was transferred from the original preservation tubes to a labelled sterile 2 mL microcentrifuge tube, then 450 µL of Chloroform:Isoamyl alcohol (ThermoFisher Scientific) was added. Tubes were vortexed and centrifuged at 15,000 × g for 5 mins, following which 400 µL of the aqueous layer was transferred to a new sterile 2 mL microcentrifuge tube, and 350 µL of ice-cold isopropanol (Fisher Scientific) and 175 µL of room temperature 5 M NaCl (ThermoFisher Scientific) were added. Samples were left to precipitate at -20°C overnight, then centrifuged at 15,000 × g for 10 mins to pellet the DNA. Liquid was decanted off before 150 µL of room temperature 70% v/v EtOH was added to wash the pellet and samples were centrifuged at 15,000 × g for 5 mins. After EtOH was decanted off, a second 70% v/v EtOH wash was used as above. After EtOH was decanted off, tubes were inverted and placed on a paper towel for at least 10 mins to remove any excess liquid. Samples were dried in a vacufuge at 45°C for 15 mins, followed by air drying until no visible EtOH remained. Finally, DNA was rehydrated with 100 µL of 1x TE buffer (ThermoFisher Scientific). Buffer-only extraction blanks were included for each batch of extractions ($n$ = 23). DNA extracts were stored at −20°C until quantification and plating.

*Microfluidic eDNA metabarcoding*

Microfluidic eDNA metabarcoding, including PCR amplification, Illumina sequencing, and demultiplexing using QIIME 2™ (Bolyen et al., 2019), was conducted by the Roy J. Carver Biotechnology Center Functional Genomic Unit at the UIUC. The Fluidigm 48.48 Access Array™ (Fluidigm Corporation, 2016) was used for PCR amplification of eDNA samples with our validated primer panel. The Fluidigm 48.48 Access Array™ uses integrated fluidic circuits and a 4-primer amplicon tagging scheme in which target-specific primer pairs amplify up to 48 different targets, allowing for the simultaneous amplification of barcoded targets in each of 2,304 reaction chambers. Forward and reverse primers were modified at the 5' end by the addition of common sequence tags (CS1 and CS2, www.fluidigm.com) which serve as the binding site for P5 and P7 Illumina sequences and dual-index multiplex barcodes. Each sample (2 ng) was amplified once with the primer panel using the Fluidigm 48.48 Access Array™. Products were quantified on a Qubit™ Fluorometer and stored at -20°C.

All samples were run on a Fragment Analyzer (Advanced Analytics, Ames, IA) and amplicon regions and expected sizes were confirmed. Samples were then pooled in equal amounts according to product concentration. The pooled products were size selected on a 2% agarose E-gel (Life Technologies) and extracted from the isolated gel slice with a QIAquick Gel Extraction Kit (Qiagen). Purified products were run on an Agilent Bioanalyzer to confirm appropriate amplicon profile and determination of average size. The products were quantified using a Qubit™ Fluorometer, pooled evenly, and the final libraries were quantified by qPCR on a BioRad CFX Connect Real-Time System (Bio-Rad Laboratories, Inc. CA). The libraries were denatured, spiked with 20% PhiX Control v3, and loaded at 8 pM to an Illumina MiSeq for cluster formation and sequencing with MiSeq Reagent Kit v3 (600-cycle) (Illumina, Inc.). In total, 40 baseline controls, 336 eDNA samples, 18 field blanks, 18 extraction blanks, two PCR negative controls, and 18 PCR positive controls were sequenced across two runs. Sequences were sorted and demultiplexed by primer set and sample, and quality and yield assessed using the FastX Tool Kit.

*Bioinformatic processing*

The demultiplexed FASTQ files for each primer set were processed using metaBEAT v0.97.11 (https://github.com/HullUnibioinformatics/metaBEAT) which incorporates open-source software for bioinformatic processing. Quality filtering (phred score Q30) and trimming were performed using Trimmomatic v0.32 (Bolger et al., 2014), merging (10 bp overlap minimum and 10% mismatch maximum) was conducted with FLASH v1.2.11 (Magoč & Salzberg, 2011), chimeras were detected using the uchime algorithm (Edgar et al., 2011) in vsearch v1.1 (Rognes et al., 2016), and clustering (97% identity with three sequences minimum per cluster) was performed with vsearch v1.1 (Rognes et al., 2016). Non-redundant query sequences were compared against our reference database using BLAST (Zhang et al., 2000). Putative taxonomic identity was assigned using a lowest common ancestor (LCA) approach based on the top 10% BLAST matches for any query matching with at least 90% identity to a reference sequence across more than 80% of its length. Unassigned sequences were subjected to a separate BLAST against the complete NCBI nucleotide (nt) database at 90% identity to determine the source via LCA as described above.

In addition to metaBEAT, the demultiplexed FASTQ files for each primer set were processed using the Anacapa Toolkit v1 (Curd et al., 2019), which has been deposited on GitHub (https://github.com/limey-bean/Anacapa) and permanently archived (http://doi.org/10.5281/zenodo.3064152). Custom reference sequence databases were generated using the module Creating Reference libraries Using eXisting tools (CRUX) which performed *in silico* amplification using ecoPCR on the EMBL standard nucleotide database. The seed database was then queried (16 November 2019) against the NCBI nt database using blastn (Camacho et al., 2009), retaining only the longest version of each sequence then retrieving taxonomy using entrez-qimme v2.0 (Baker, 2016). CRUX generated an unfiltered database with all accessions and taxonomic path information, a filtered database that excludes accessions with ambiguous taxonomic paths, and a Bowtie2-formated index library (Langmead & Salzberg, 2012). The second module conducted DNA sequence quality control and amplicon sequence variant (ASV) parsing using DADA2 v1.14 (Callahan et al., 2016). First, cutadapt (Martin, 2011) and FastX-toolkit (Gordon & Hannon, 2010) were used to trim primers, Illumina adapters, and low-quality bases from raw FASTQ files for each sample using the same quality filtering and trimming settings as in metaBEAT. A custom Python script sorted sequence reads into three sets – paired-end, forward only, and reverse only – then processed each set separately with DADA2 to denoise, dereplicate, merge paired reads, and remove chimeric sequences. This module generated ASV FASTA files and ASV count summary tables for four read types: merged paired-end reads, unmerged paired-end reads, forward only reads, and reverse only reads. The resulting ASV FASTA files and count summary tables were inputted into the Anacapa Classifier module, which assigns taxonomy to ASVs using Bowtie2 and a modified version of Bayesian Least Common Ancestor (BLCA) algorithm (Gao et al., 2017). Bowtie2 queried ASVs against the CRUX-generated reference databases using the very-sensitive option to determine up to 100 reference matches. Bowtie2-BCLA was used to process the output, using muscle v3.8.31 (Edgar, 2004) for multiple sequence alignment with 100 bootstraps to probabilistically determine taxonomic identity by selecting the lowest common ancestor from the multiple weighted Bowtie2 hits for each ASV (Curd et al., 2019).

metaBEAT and Anacapa use a different suite of open-source software for bioinformatic processing, which allowed us to assess the robustness of eDNA detections. metaBEAT produced comparable or better detection than Anacapa for our focal pollinator species and species known to occur in the greenhouse (Figure S1), thus we used these results for downstream analyses. metaBEAT and Anacapa shared 30 metazoan taxa after application of a false positive sequence threshold specific to each pipeline. Unique species of note detected with Anacapa were *Euhybus* *triplex*, eastern flower thrip (*Frankliniella tritici*), white-lined sphinx (*Hyles lineata*), *Liposcelis decolor*, insidious flower bug (*Orius insidiosus*), *Orthoperus scutellaris*, and *Philygria debilis* (Figure S1). However, metaBEAT detected the Asian tiger mosquito (*Aedes albopictus*), *Amiota setigera*, banded garden spider (*Argiope trifasciata*), silverleaf whitefly (*Bemisia tabaci*), fungus knat (*Bradysia impatiens*), *Elachodelphax metcalfi*, *Homidia socia*, purple-scum springtail (*Hypogastrura vernalis*), *Incertella bispina*, *Islandiana flaveola*, cigarette beetle (*Lasioderma serricorne*), *Leucopis piniperda*, *Malloewia setulosa*, *Meioneta unimaculata*, *Melanophthalma inermis*, housefly (*Musca domestica*), *Psammotettix lividellus*, *Scaptomyza pallida*, *Scatella stagnalis*, *Simulium vittatum*, *Tanytarsus dendyi*, *Telenomus podisi*, firebrat (*Thermobia domestica*), eastern calligrapher (*Toxomerus geminatus*), and *Trachyopella nuda*.

#### Tables

**Table S1.** Flower species that comprised the greenhouse flower assemblage. These flowering plants provided a constant supply of pollen and nectar to the common eastern bumblebee (*Bombus impatiens*) colony introduced to the greenhouse.

| **Binomial name** | **Common name(s)** |
|---|---|
| *Ammi majus* | Queen Anne's Lace |
| *Calendula officinalis* | Pot Marigold |
| *Centaurea cyannus* | Cornflower |
| *Clarkia amoena* | Farwell-To-Spring, Godetia |
| *Clarkia unguiculata* | Elegant Clarkia, Mountain Garland |
| *Coreopsis tinctoria* | Plains Coreopsis |
| *Cosmos bipinnatus* | Garden Cosmos |
| *Cosmos sulphureus* | Sulphur Cosmos |
| *Cynoglossum amabile* | Chinese Forget-Me-Not |
| *Delphinium ajacis* | Rocket Larkspur |
| *Eschscholzia californica* | California Poppy |
| *Gaillardia pulchella* | India Garland |
| *Gilia capitata* | Globe Gillia, Blue-Thimble-Flower |
| *Gypsophila elegans* | Baby's Breath |
| *Helianthus annuus* | Dwarf Sunflower Sunspot |
| *Lavatera trimestris* | Rose Mallow |
| *Linaria maroccana* | Baby Snapdragon, Moroccan Toadflax |
| *Linum grandiflorum "Rubrum"* | Scarlet Flax |
| *Lupinus* | Lupine |
| *Mirabilis jalapa* | Four O'Clock |
| *Nemophila menziesii* | Baby Blue Eyes |
| *Papaver rhoeas* | Red Poppy, Common Poppy |
| *Silene armeria* | None-So-Pretty, Sweet William Catchfly |

**Table S2.** Reagent volumes that were used for ATL preservation and QBT extraction or CTAB preservation and PCI extraction of different sample types collected from our four focal flower species.

| Flower species | QBT | | | | | | PCI | | |
|---|---|---|---|---|---|---|---|---|---|
| | Whole flower | | Swab | | Nectar draw | | Whole flower | Swab | Nectar draw |
| | ATL (µL) | PK (µL) | ATL (µL) | PK (µL) | ATL (µL) | PK (µL) | CTAB (µL) | CTAB (µL) | CTAB (µL) |
| *Penstemon* | 900 | 200 | 600 | 100 | - | - | 900 | 600 | - |
| *Monarda* | 1200 | 300 | 600 | 100 | - | - | 1200 | 600 | - |
| *Solanum* | 600 | 100 | - | - | - | - | 600 | 600 | - |
| *Cynoglossum amabile* | - | - | - | - | 300 | 60 | - | - | 300 |

**Table S3.** Results of *in silico* primer testing for each primer pair considered for microfluidic eDNA metabarcoding. The percentage of species that amplified when allowing up to 3 mismatches in each primer with a custom reference database containing sequences for 5313 pollinator species native to Illinois is given.

| Primer set | Forward primer (5' – 3') | Reverse primer (5' – 3') | Marker | Reported fragment size (bp) | Target taxa | Reference | Mean fragment size (bp) | Min. fragment size (bp) | Max. fragment size (bp) | Percentage of species amplified |
|---|---|---|---|---|---|---|---|---|---|---|
| Ins16S-1F/ Ins16S-1Rshort | TRRGACGAGAAGAC CCTATA | ACGCTGTTATCCCT AARGTA | 16S | 156 | Invertebrates | Clarke et al. (2014) | 143 | 81 | 259 | 10.90 |
| 16SMAV-F/ 16SMAV-R | CCAACATCGAGGTC RYAA | ARTTACYNTAGGGA TAACAG | 16S | 36 | Invertebrates | De Barba et al. (2014) | NA | NA | NA | 0 |
| MOL16S_F/ MOL16S_R | RRWRGACRAGAAG ACCCT | ARTCCAACATCGAG GT | 16S | 183-310 | Molluscs | Klymus et al. (2017) | 202 | 109 | 321 | 10.75 |
| SPH16S_F/ SPH16S_R | TAGGGGAAGGTATG AATGGTTTG | ACATCGAGGTCGCA ACC | 16S | 299 | Sphaeriidae | Klymus et al. (2017) | 299 | 298 | 300 | 0.06 |
| BF1/ BR1 | ACWGGWTGRACWG TNTAYCC | ARYATDGTRATDGC HCCDGC | COI | 217 | Invertebrates | Elbrecht & Leese (2017) | 212 | 61 | 383 | 46.45 |
| BF1/ BR2 | ACWGGWTGRACWG TNTAYCC | TCDGGRTGNCCRAA RAAYCA | COI | 316 | Invertebrates | Elbrecht & Leese (2017) | 315 | 256 | 482 | 12.67 |
| BF2/ BR2 | GCHCCHGAYATRGC HTTYCC | TCDGGRTGNCCRAA RAAYCA | COI | 421 | Invertebrates | Elbrecht & Leese (2017) | 420 | 361 | 459 | 9.94 |
| BF3/ BR2 | CCHGAYATRGCHTT YCCHCG | TCDGGRTGNCCRAA RAAYCA | COI | 418 | Arthropods | Elbrecht et al.(2019), Elbrecht & Leese (2017) | 417 | 358 | 456 | 9.98 |

| Primer set | Forward primer (5' – 3') | Reverse primer (5' – 3') | Marker | Reported fragment size (bp) | Target taxa | Reference | Mean fragment size (bp) | Min. fragment size (bp) | Max. fragment size (bp) | Percentage of species amplified |
|---|---|---|---|---|---|---|---|---|---|---|
| nsCOIFo/ mlCOIintK | THATRATNGGNGGN TTYGGNAAHTG | GGRGGRTAWACWG TTCAWCCWGTWCC | COI | 124 | Invertebrates | Günther et al. (2018) | 123 | 88 | 130 | 44.02 |
| mlCOIintF jgHCO2198 | GGWACWGGWTGAA CWGTWTAYCCYCC | TAIACYTCIGGRTGIC CRAARAAYCA | COI | 313 | Metazoans | Leray et al. (2013), Geller et al. (2013) | 312 | 298 | 314 | 11.71 |
| fwhF1/ fwhR1 | YTCHACWAAYCAY AARGAYATYGG | ARTCARTTWCCRAA HCCHCC | COI | 178 | Invertebrates | Vamos et al. (2017) | 178 | 154 | 270 | 4.74 |
| fwhF2/ fwhR2 | GGDACWGGWTGAA CWGTWTAYCCHCC | GTRATWGCHCCDGC AARWACWGG | COI | 205 | Invertebrates | Vamos et al. (2017) | 204 | 145 | 219 | 46.09 |
| fwhF2/ fwhR22n | GGDACWGGWTGAA CWGTWTAYCCHCC | GTRATWGCHCCDGC TARWACWGG | COI | 205 | Invertebrates | Vamos et al. (2017) | 204 | 145 | 219 | 46.23 |
| Uni-MinibarF1/ Uni-MinibarR1 | TCCACTAATCACAA RGATATTGGTAC | GAAAATCATAATGA AGGCATGAGC | COI | 127 | Metazoans | Meusnier et al. (2008) | 127 | 127 | 127 | 0.17 |
| Ill-B-F/ Ill-HCO2198 | CCIGAYATRGCITTY CCICG | TAAACTTCAGGGTG ACCAAAAAATCA | COI | Not reported | Invertebrates | Shokralla et al. (2015) | 417 | 403 | 419 | 6.64 |
| Ill-LCO1490/ Ill-C-R | GGTCAACAAATCAT AAAGATATTGG | GGIGGRTAIACIGTTC AICC | COI | Not reported | Invertebrates | Shokralla et al. (2015) | NA | NA | NA | 0 |
| MG-LCO1490-MiSeq/MG-R-MiSeq (LepF1/ EPT-long-univR) | ATTCHACDAAYCAY AARGAYATYGG | ACTATAAAARAAAA TYTDAYAAADGCRT | COI | 133 | Arthropods | Galan et al. (2018), Hajibabaei et al.(2011), Hebert et al. (2004) | 133 | 109 | 321 | 4.59 |
| ZBJ-ArtF1c/ZBJ-ArtR2c | AGATATTGGAACWT TATATTTTATTTTTG G | WACTAATCAATTW CCAAATCCTCC | COI | 157 | Arthropods | Zeale et al. (2011) | 157 | 133 | 217 | 4.40 |

| Primer set | Forward primer (5' – 3') | Reverse primer (5' – 3') | Marker | Reported fragment size (bp) | Target taxa | Reference | Mean fragment size (bp) | Min. fragment size (bp) | Max. fragment size (bp) | Percentage of species amplified |
|---|---|---|---|---|---|---|---|---|---|---|
| LepLCO/ McoiR1 | RKTCAACMAATCAT AAAGATATTGG | AATCCBCCRATTAW AATKGGTAT | COI | ~150 | Invertebrates | Corse et al. (2019) | 162 | 139 | 163 | 3.33 |
| LFCR (LepLCO/McoiR2) | RKTCAACMAATCAT AAAGATATTGG | CCBCCRATTAWAAT KGGTATHAC | COI | ~150 | Invertebrates | Corse et al. (2019) | 159 | 136 | 160 | 3.20 |
| LepLCO / MLepF1rev | RKTCAACMAATCAT AAAGATATTGG | CGTGGAAAWGCTAT ATCWGGTG | COI | ~150 | Invertebrates | Corse et al. (2019) | 217 | 194 | 218 | 2.99 |
| MFZR (Uni-MinibarF1/ ZBJ-ArtR2c) | TCCACTAATCACAA RGATATTGGTAC | WACTAATCAATTW CCAAATCCTCC | COI | ~230 | Invertebrates | Corse et al. (2017) Meusnier et al. (2008), Zeale et al. (2011) | 175 | 175 | 175 | 1.19 |

**Table S4.** Summary of bioinformatic processing with metaBEAT for each primer set used for microfluidic eDNA metabarcoding. For all primer sets, quality filtering used phred score Q30, paired and forward only reads were retained after merging, chimera detection was performed using vsearch with the uchime algorithm, clustering was performed at 97% identity with a minimum of three query sequences per cluster, and query sequences had to match with at least 90% identity to a reference sequence across more than 80% of its length for BLAST. Two sets of read counts are given for each primer set as two Illumina MiSeq runs were required to sequence all samples. The number of taxonomically assigned reads and unassigned reads after bioinformatic processing are given as well as the number and percentage of taxonomically assigned reads that belonged to Arthropoda and Apidae.

| **Primer set** | **Raw sequence reads** | **Read crop** | **Minimum read length** | **PCR product length (bp) ± deviation** | **Taxonomically assigned reads** | **Reads assigned to Arthropoda** | **Reads assigned to Apidae** | **Unassigned reads** |
|---|---|---|---|---|---|---|---|---|
| BF1/BR1 | 223,252 | 217 | 200 | 217 ± 10% | 82,750 | 81,069 (97.97%) | 1,006 (1.22%) | 14,611 |
| | 134,826 | | | | 56,092 | 53,562 (95.49%) | 1,217 (2.17%) | 1,643 |
| BF1/BR2 | 834,984 | 200 | 200 | 316 ± 10% | 348,426 | 345,082 (99.04%) | 4,284 (1.23%) | 13,795 |
| | 747,384 | | | | 299,612 | 271,635 (90.66%) | 11,136 (3.72%) | 19,492 |
| BF2/BR2 | 927,980 | 250 | 200 | 421 ± 10% | 272,724 | 193,056 (70.79%) | 1,308 (0.48%) | 37,421 |
| | 746,132 | | | | 159,547 | 128,337 (80.44%) | 1,158 (0.73%) | 98,432 |
| BF3/BR2 | 745,552 | 250 | 200 | 418 ± 10% | 219,469 | 145,255 (66.19%) | 2,004 (0.91%) | 34,601 |
| | 567,318 | | | | 133,530 | 108,673 (81.39%) | 1,313 (0.98%) | 64,692 |
| fwhF1/fwhR1 | 484,574 | 178 | 150 | 178 ± 10% | 201,390 | 143,632 (71.32%) | 122 (0.06%) | 6,230 |
| | 440,484 | | | | 188,062 | 186,913 (99.39%) | 919 (0.49%) | 472 |
| fwhF2/fwhR2n | 1,958,032 | 205 | 190 | 205 ± 10% | 827,997 | 798,016 (96.38%) | 24,124 (2.91%) | 18,042 |
| | 1,842,304 | | | | 756,621 | 745,802 (98.57%) | 60,092 (7.94%) | 15,925 |
| LepLCO/McoiR1 | 430,772 | 163 | 100 | 163 ± 10% | 138,925 | 138,467 (99.67%) | 0 (0.00%) | 51,856 |
| | 372,830 | | | | 165,098 | 152,498 (92.37% | 23 (0.01%) | 6 |
| LepLCO/McoiR2 | 98,710 | 160 | 100 | 160 ± 10% | 41,950 | 41,950 (100.00%) | 0 (0.00%) | 12 |
| | 100,234 | | | | 38,672 | 38,672 (100.00%) | 210 (0.54%) | 2,977 |
| LepLCO/MLepF1rev | 668,790 | 218 | 200 | 218 ± 10% | 284,017 | 281,842 (99.23%) | 7,377 (2.60%) | 4,355 |
| | 856,360 | | | | 362,698 | 362,698 (100.00%) | 7,801 (2.15%) | 3 |
| mlCOIintF/jgHCO2198 | 3,844,444 | 200 | 200 | 313 ± 10% | 1,522,791 | 1,483,344 (97.41%) | 30,207 (1.98%) | 83,545 |
| | 3,516,470 | | | | 1,331,270 | 1,279,376 (96.10%) | 47,861 (3.60%) | 126,281 |
| nsCOIFo/mlCOIintK | 443,262 | 124 | 100 | 124 ± 10% | 54,546 | 54,381 (99.70%) | 1,096 (2.01%) | 81.484 |
| | 234,228 | | | | 26,209 | 26,168 (99.84%) | 3,697 (14.11%) | 45,703 |
| Uni-MinibarF1/Uni-MinibarR1 | 6,824 | 127 | 100 | 127 ± 10% | 2,568 | 1,041 (40.54%) | 0 (0.00%) | 301 |
| | 4,556 | | | | 1,814 | 1,808 (99.67%) | 4 (0.22%) | 129 |
| ZBJ-ArtF1c/ZBJ-ArtR2c | 2,290,880 | 157 | 100 | 157 ± 10% | 992,475 | 992,059 (99.96%) | 482 (0.05%) | 40,737 |
| | 1,635,376 | | | | 731,397 | 731,397 (100.00%) | 10 (0.001%) | 1,120 |
| Ins16S-1F/Ins16S-1Rshort | 4,743,478 | 155 | 80 | 155 ± 10% | 235,479 | 207,950 (88.31%) | 3,393 (1.44%) | 1,433,913 |
| | 5,848,630 | | | | 232,577 | 232,048 (99.77%) | 1,027 (0.44%) | 1,851,875 |
| MOL16S_F/MOL16S_R | 3,069,976 | 221 | 180 | 221 ± 10% | 744,689 | 197,473 (26.52%) | 0 (0.00%) | 555,901 |
| | 2,069,098 | | | | 276,152 | 70,786 (25.63%) | 3 (0.001%) | 564,032 |

**Table S5.** Summary of tests used to assess normality of data and model residuals in analyses of alpha diversity (taxon richness) according to each grouping variable. The response variable was visually examined using histograms and quantile-quantile plots before being tested using Shapiro-Wilk Test. Variance in the response variable was compared between groups within each grouping variable using Levene's Test. A one-way Analysis of Variance (ANOVA) was performed for the response variable and each grouping variable regardless of whether the data was normally distributed and possessed equal variance. The standardised Pearson residuals from each ANOVA were examined (see Figures S9, S10 and S11) to determine whether the model assumptions were met. In all cases, the residuals vastly deviated from the model assumptions. Therefore, non-parametric tests reported in the main text were used instead.

| **Grouping variable** | **Shapiro-Wilk Test on response variable** | **Levene's Test on response variable** | **Shapiro-Wilk Test on standardised residuals (ANOVA)** |
|---|---|---|---|
| Flower species | W = 0.754<br>**P < 0.001** | F = 3.933<br>**P = 0.009** | W = 0.825<br>**P < 0.001** |
| Sample type | W = 0.754<br>**P < 0.001** | F = 2.783<br>P = 0.064 | W = 0.856<br>**P < 0.001** |
| Preservation/extraction | W = 0.754<br>**P < 0.001** | F = 0.231<br>P = 0.632 | W = 0.857<br>**P < 0.001** |

### Figures

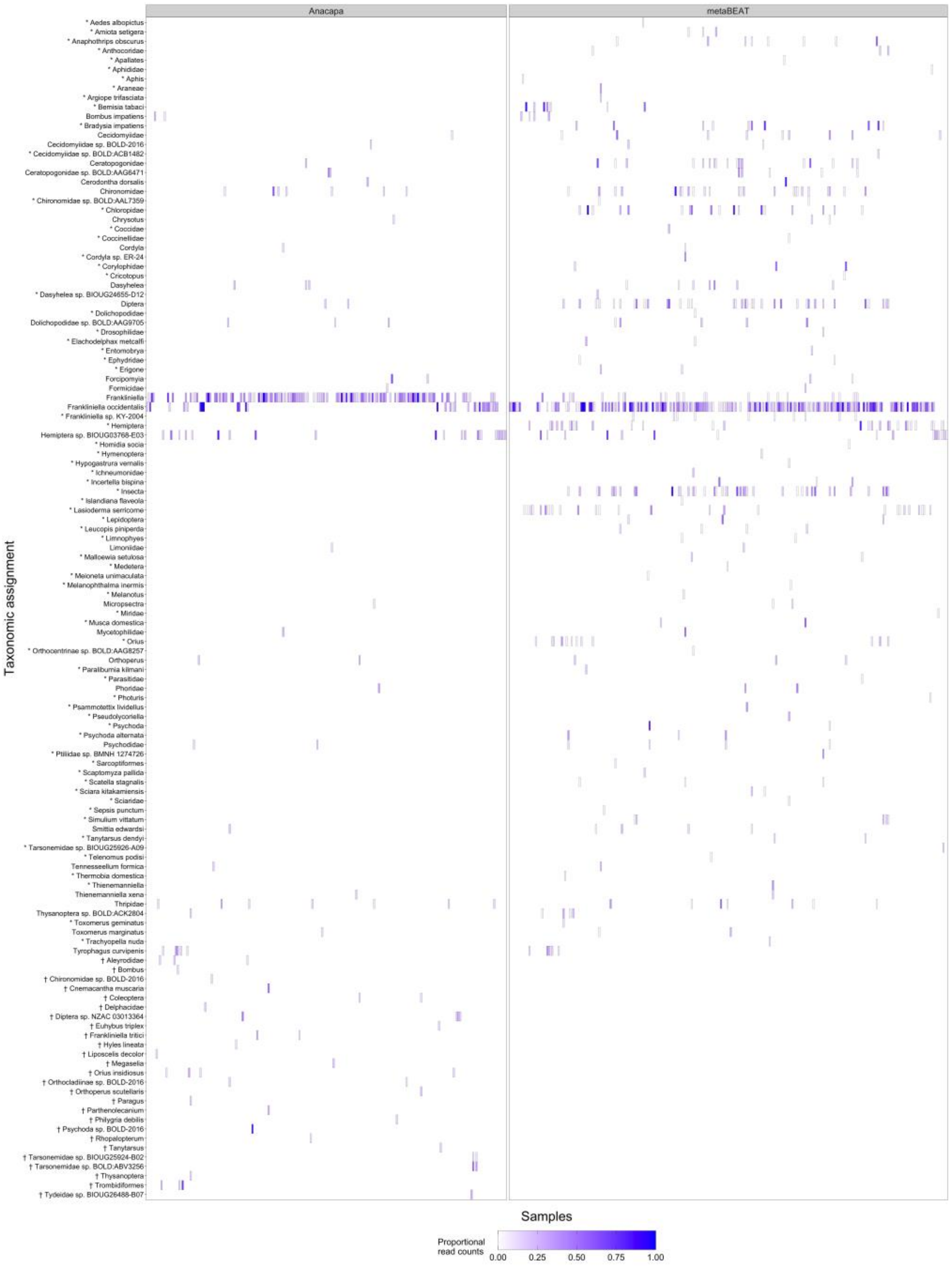


**Figure S1.** Heatmap showing proportional read counts for taxa detected in eDNA samples and baseline controls by Anacapa and metaBEAT after false positive sequence threshold application (8.02% and 2.05% respectively). Symbols after taxon names indicate taxa detected only by Anacapa (†, 26 taxa) or metaBEAT (*, 70 taxa). Taxon names with no symbol indicate taxa detected by both pipelines (30 taxa). Taxa that were not detected in a sample by each pipeline are coloured white with no border.

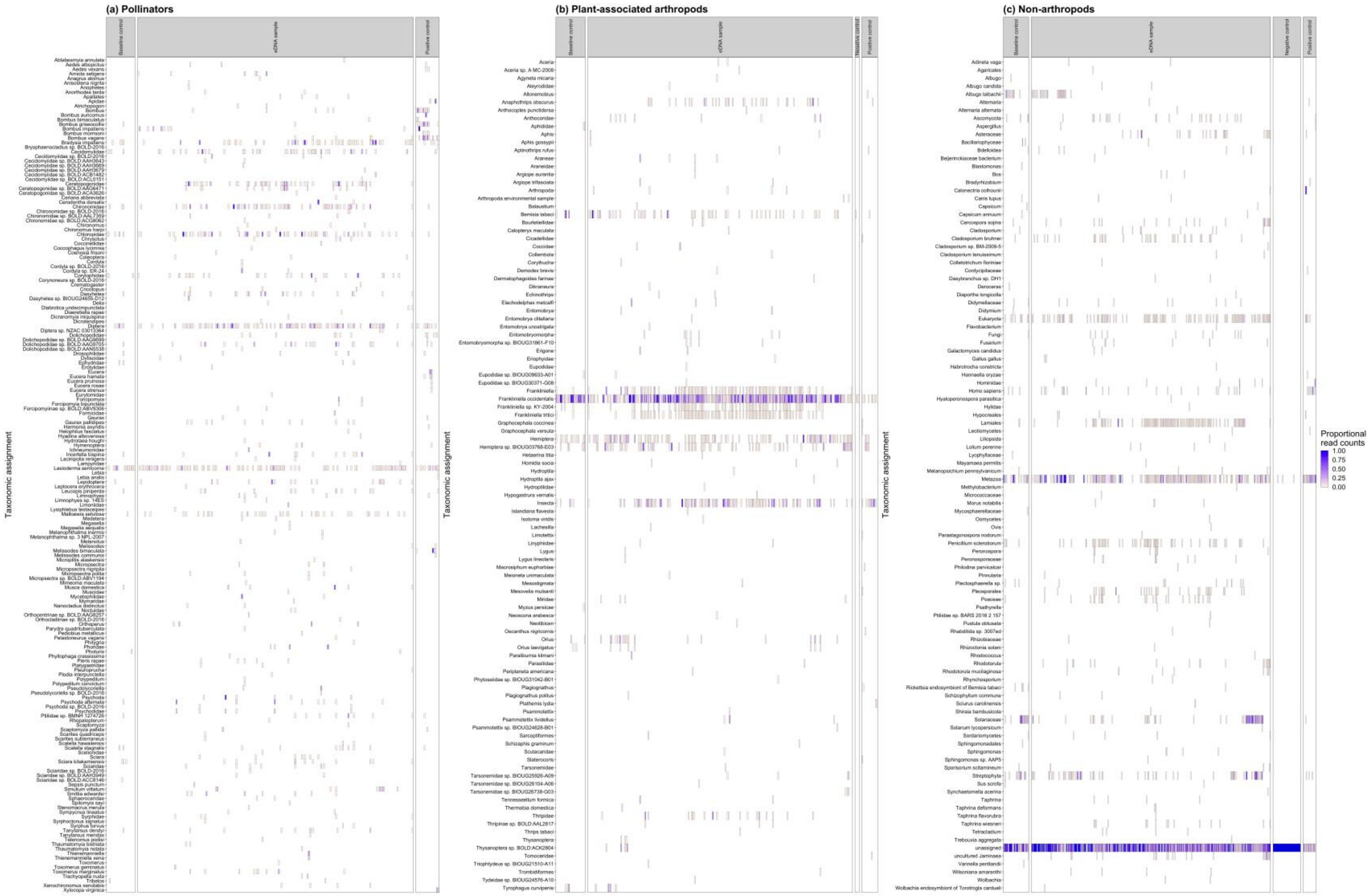


**Figure S2.** Heatmap showing proportional read counts for **(a)** pollinator, **(b)** plant-associated, and **(c)** non-arthropod taxa detected in eDNA samples, baseline controls, negative process controls, and PCR positive controls by metaBEAT before false positive sequence threshold application. Taxa that were not detected in a sample by each pipeline are coloured white with no border.

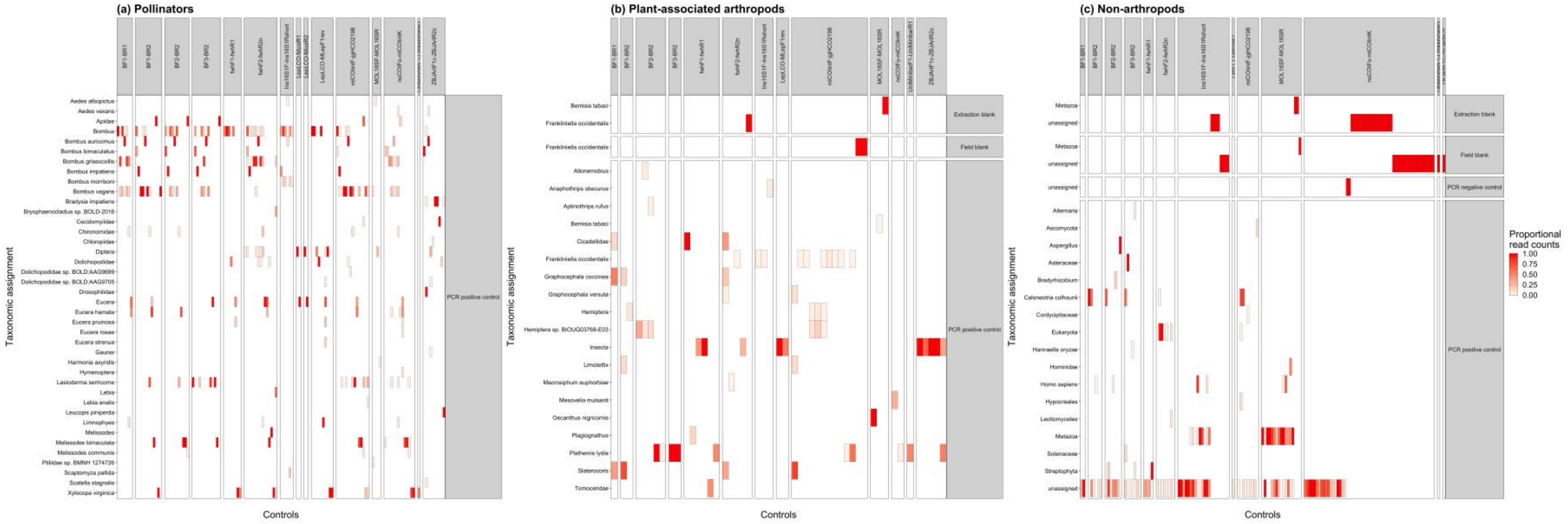


**Figure S3.** Heatmap showing proportional read counts for **(a)** pollinator, **(b)** plant-associated, and **(c)** non-arthropod taxa detected in negative process controls and PCR positive controls with different primer sets using metaBEAT before false positive sequence threshold application. Taxa that were not detected in a sample by each pipeline are coloured white with no border.

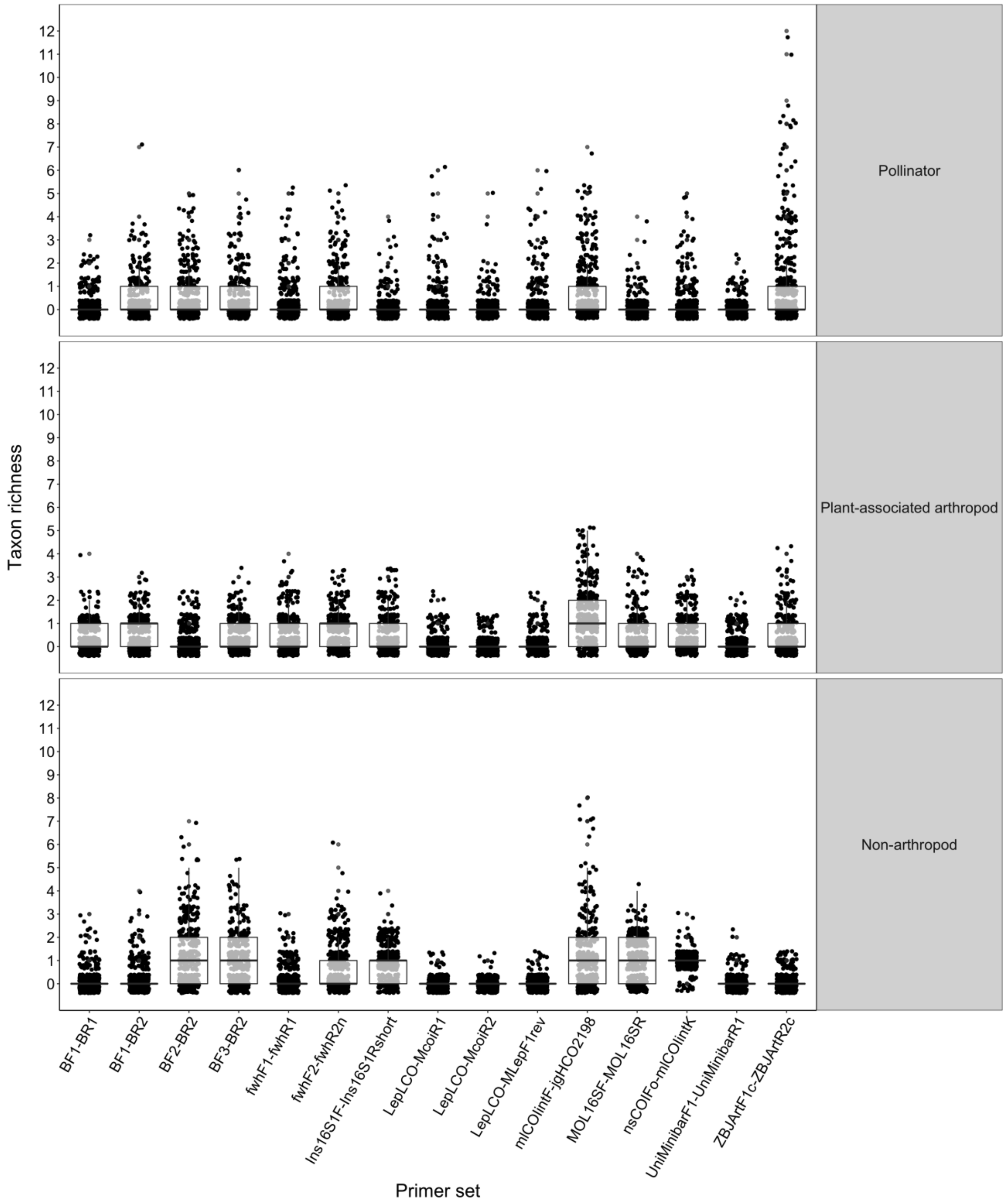


**Figure S4.** Boxplot showing the number of pollinator, plant-associated arthropod and non-arthropod taxa detected in each sample with different primer sets using metaBEAT before false positive sequence threshold application.

#### (a) Pollinators

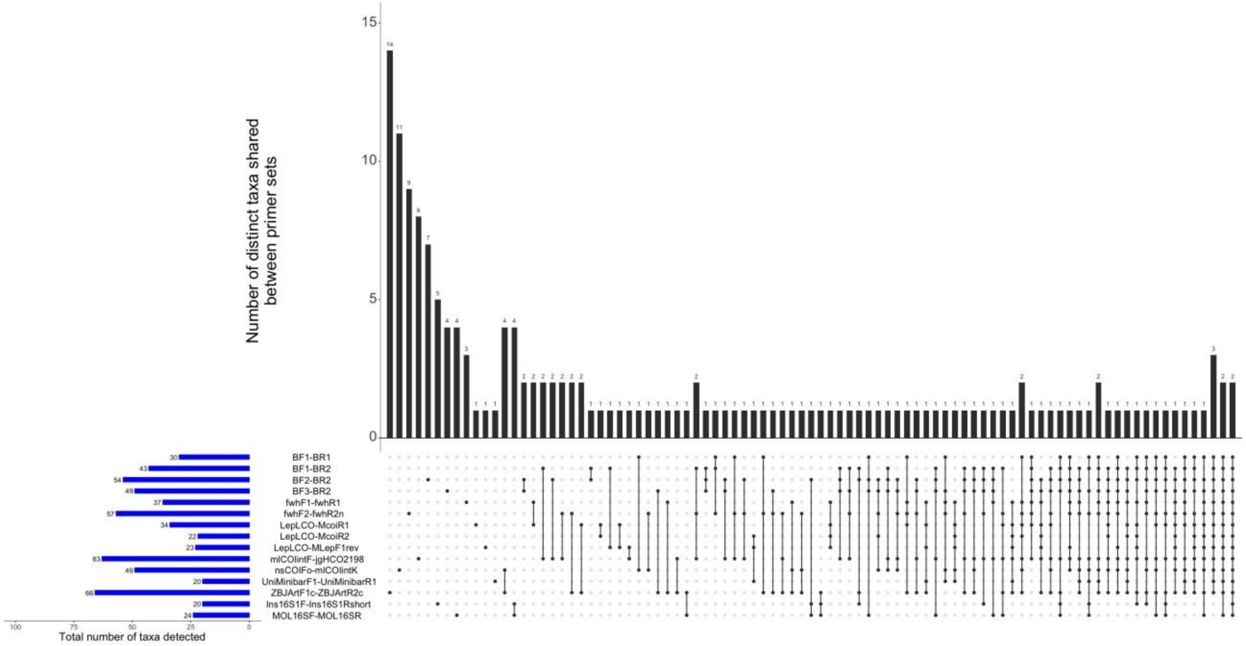


#### (b) Plant-associated arthropods

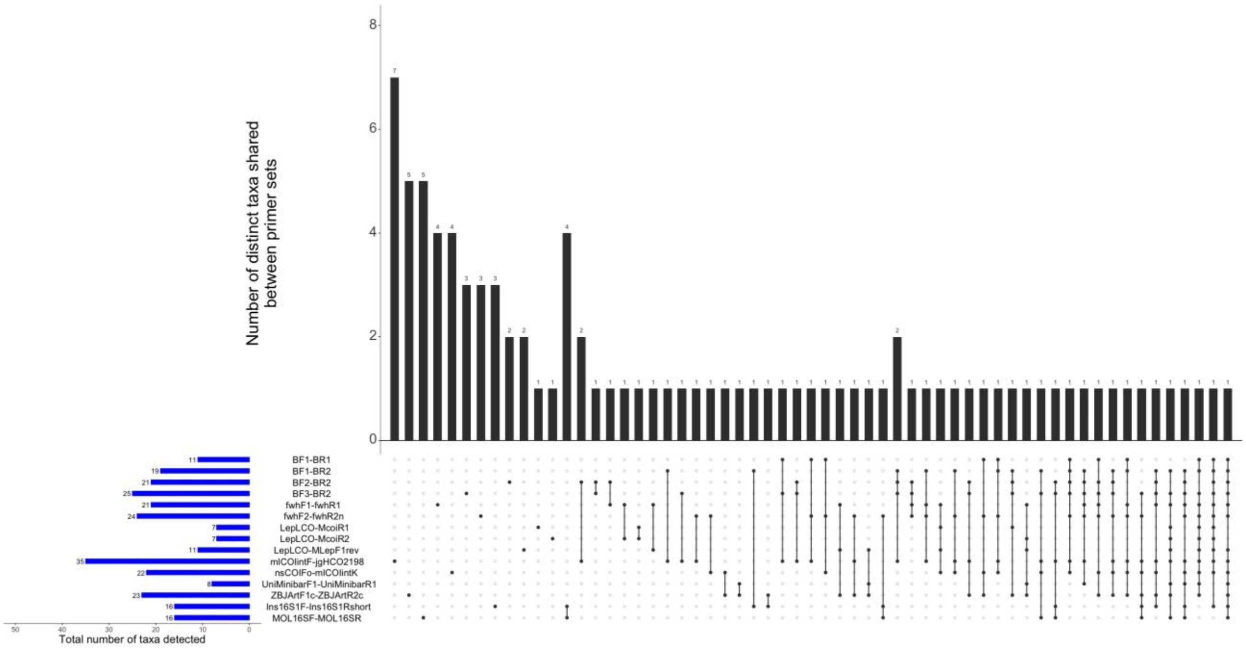

#### (c) Non-arthropods

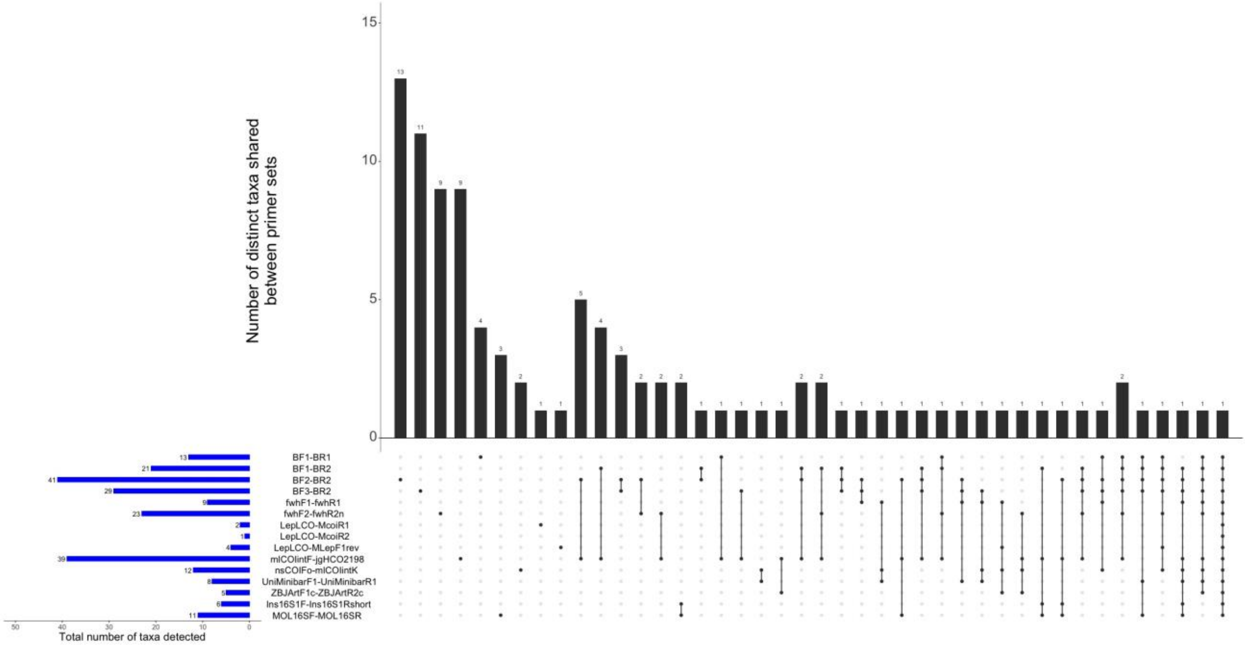


**Figure S5.** Upset plot, produced using UpSetR v1.4.0 (Gehlenborg, 2019), showing the number of distinct **(a)** pollinator, **(b)** plant-associated arthropod, and **(c)** non-arthropod taxa detected by individual primer sets and combinations of primer sets using metaBEAT before false positive threshold application. The inset plot shows the total number of taxa detected across all samples by individual primer sets.

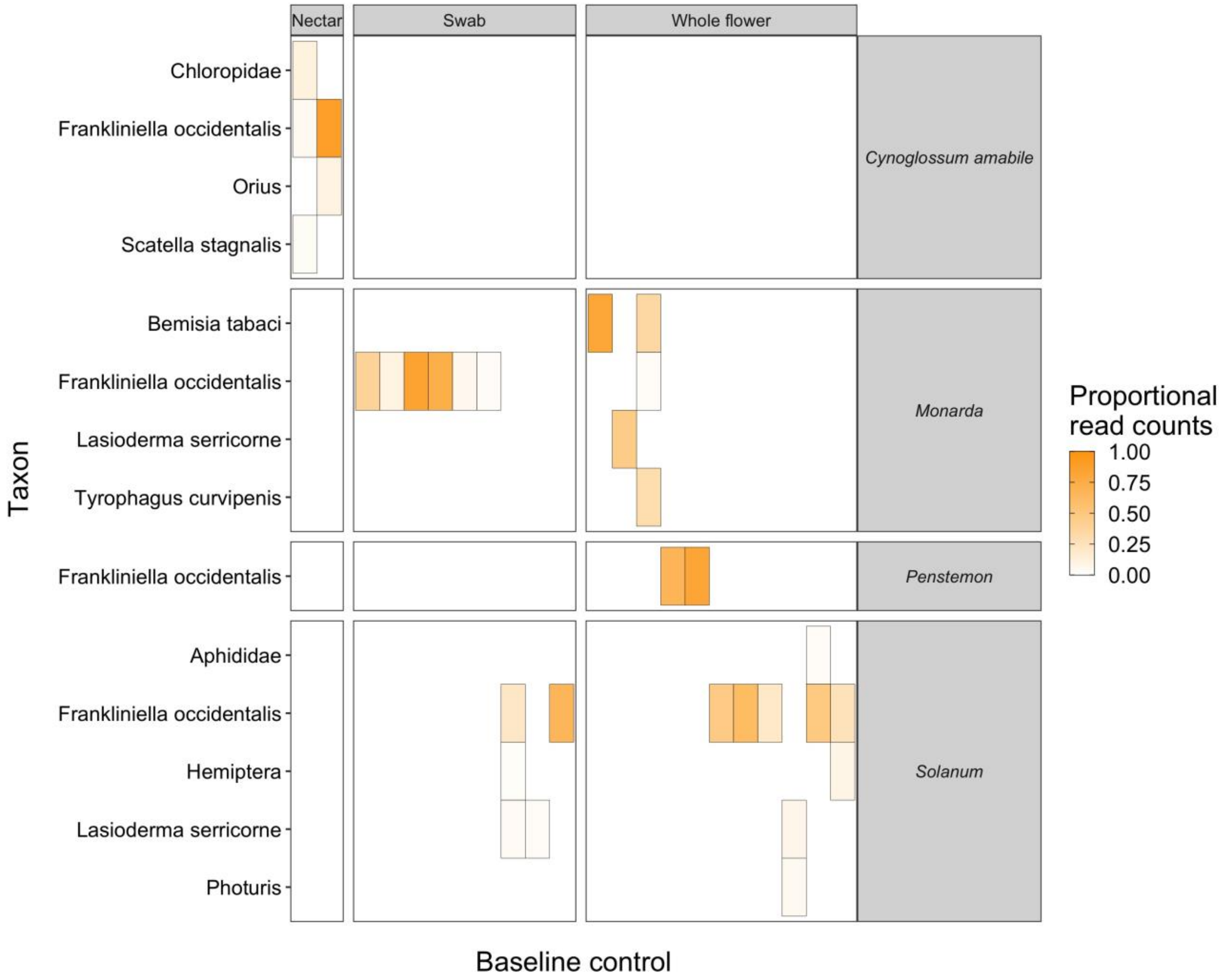


**Figure S6.** Heatmap showing the frequency at which taxa were detected in baseline controls collected from flower species before they were introduced to the secure room containing common eastern bumblebees (*Bombus impatiens*). Taxa that were not detected in a control are coloured white with no border.

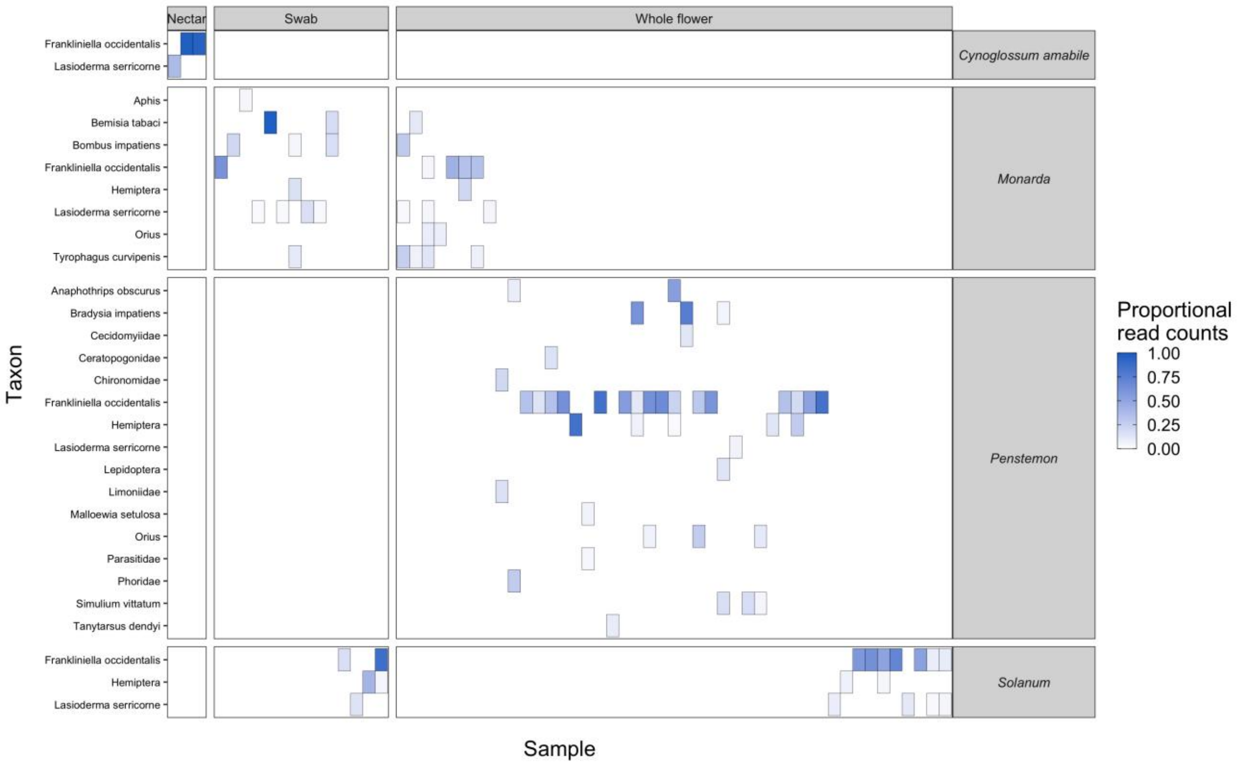


**Figure S7.** Heatmap showing the frequency at which taxa were detected in samples collected from our four focal flower species after introduction to the secure room containing common eastern bumblebees (*Bombus impatiens*). Taxa that were not detected in an eDNA sample are coloured white with no border.

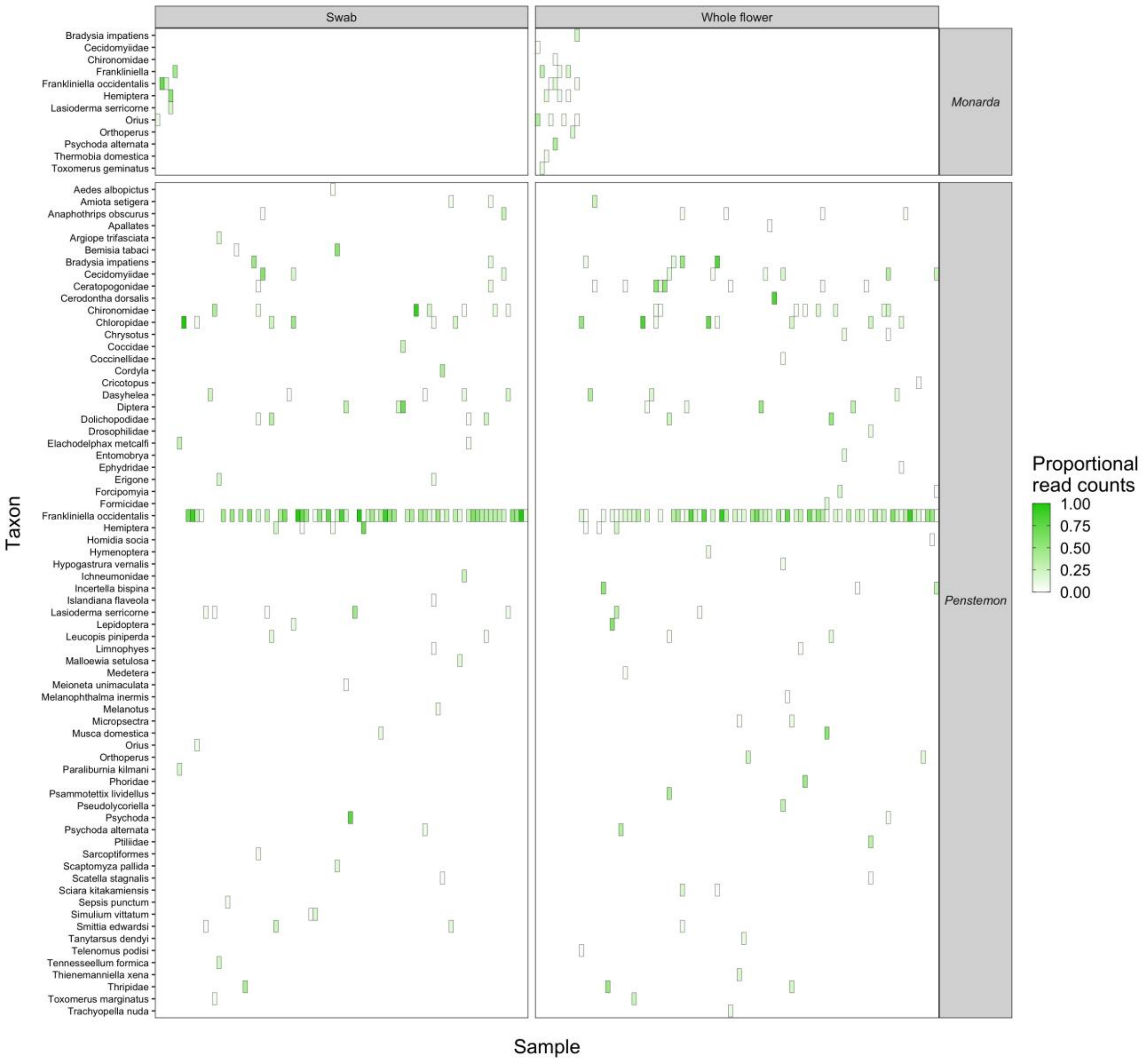


**Figure S8.** Heatmap showing the frequency at which taxa were detected in field samples collected from two focal flower species that occur on the University of Illinois at Urbana-Champaign campus. Taxa that were not detected in an eDNA sample are coloured white with no border.

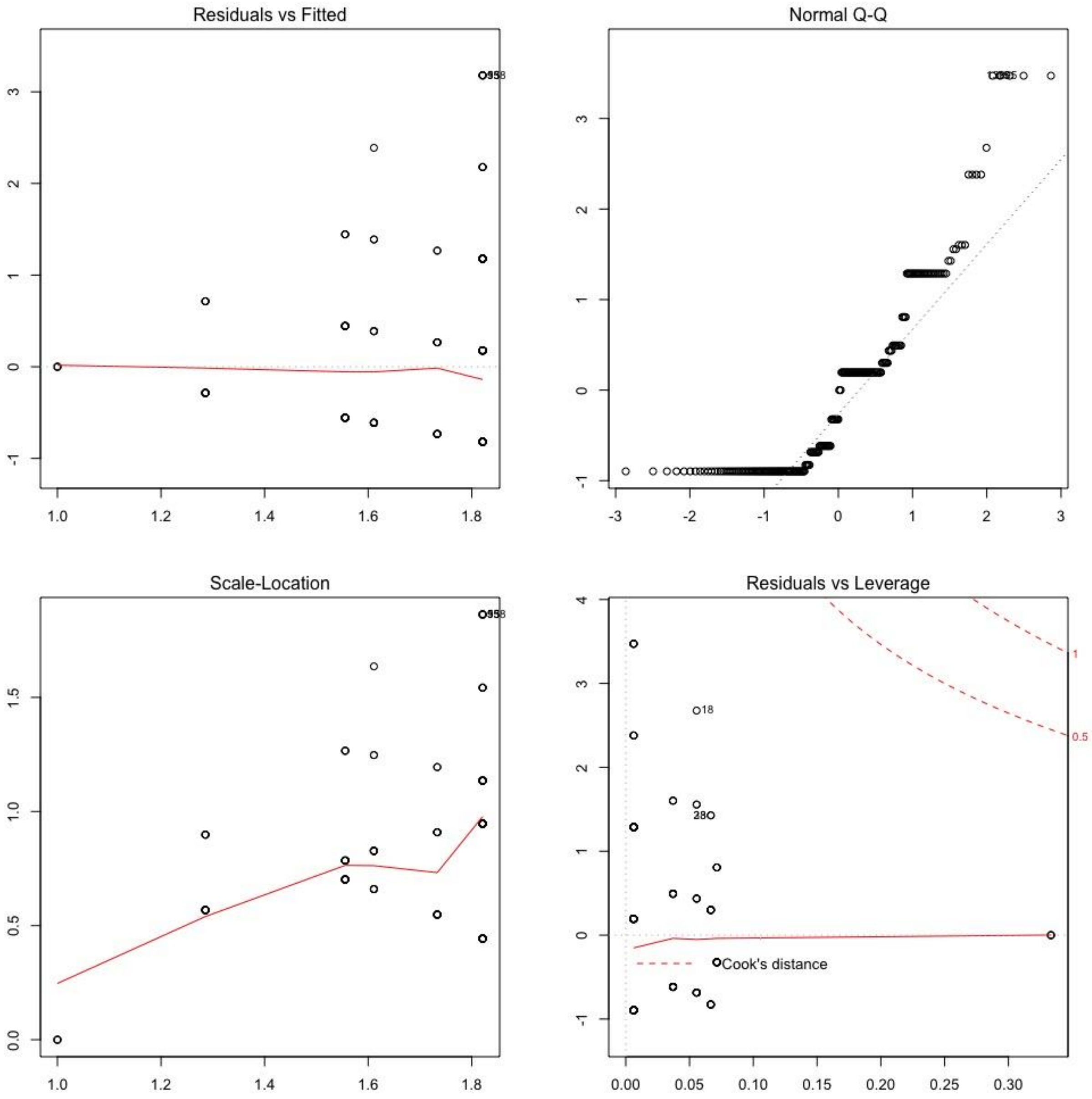


**Figure S9.** Standardised Pearson residuals from the one-way ANOVA comparing taxon richness between different flower species.

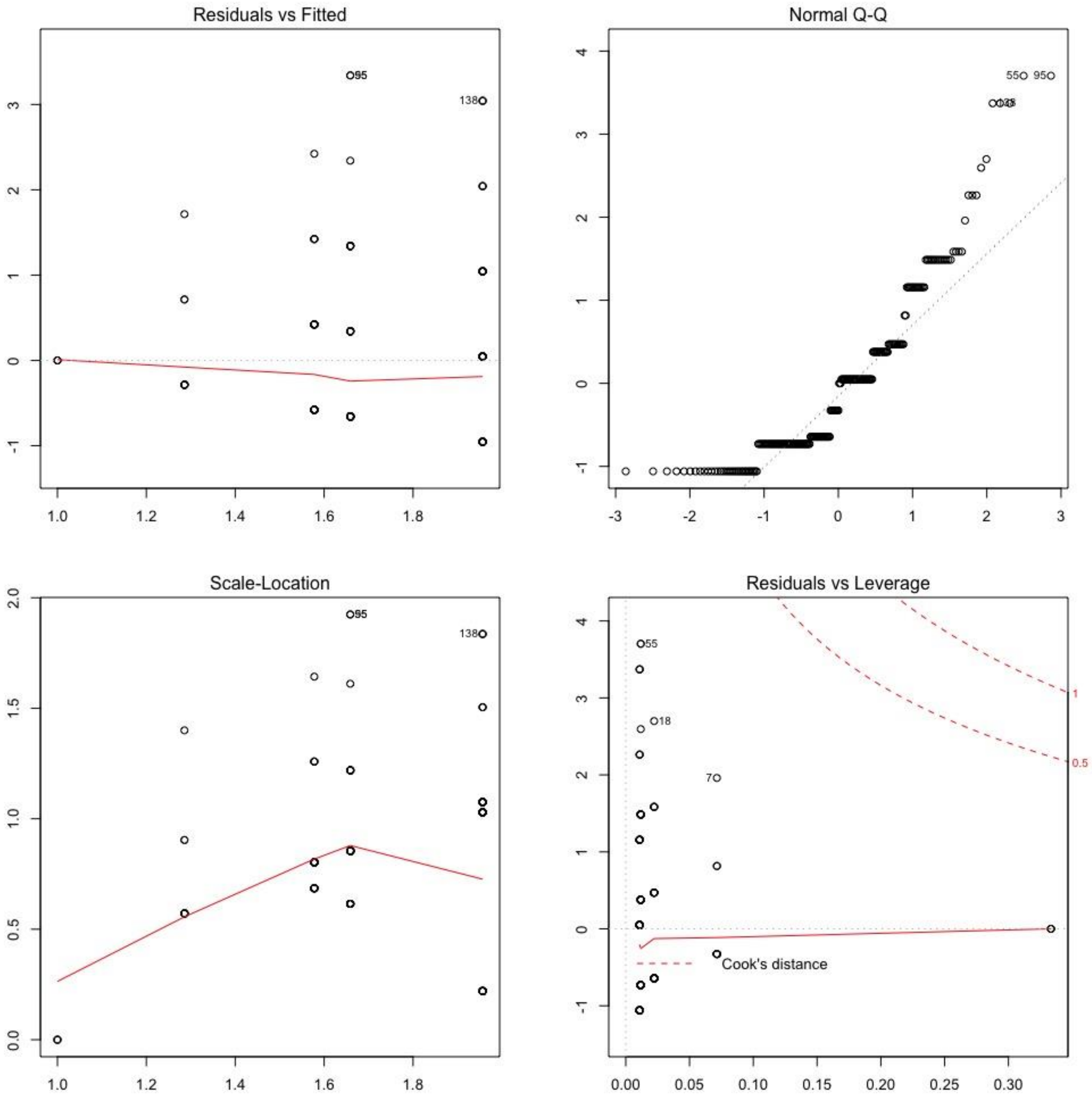


**Figure S10.** Standardised Pearson residuals from the one-way ANOVA comparing taxon richness between different sample types.

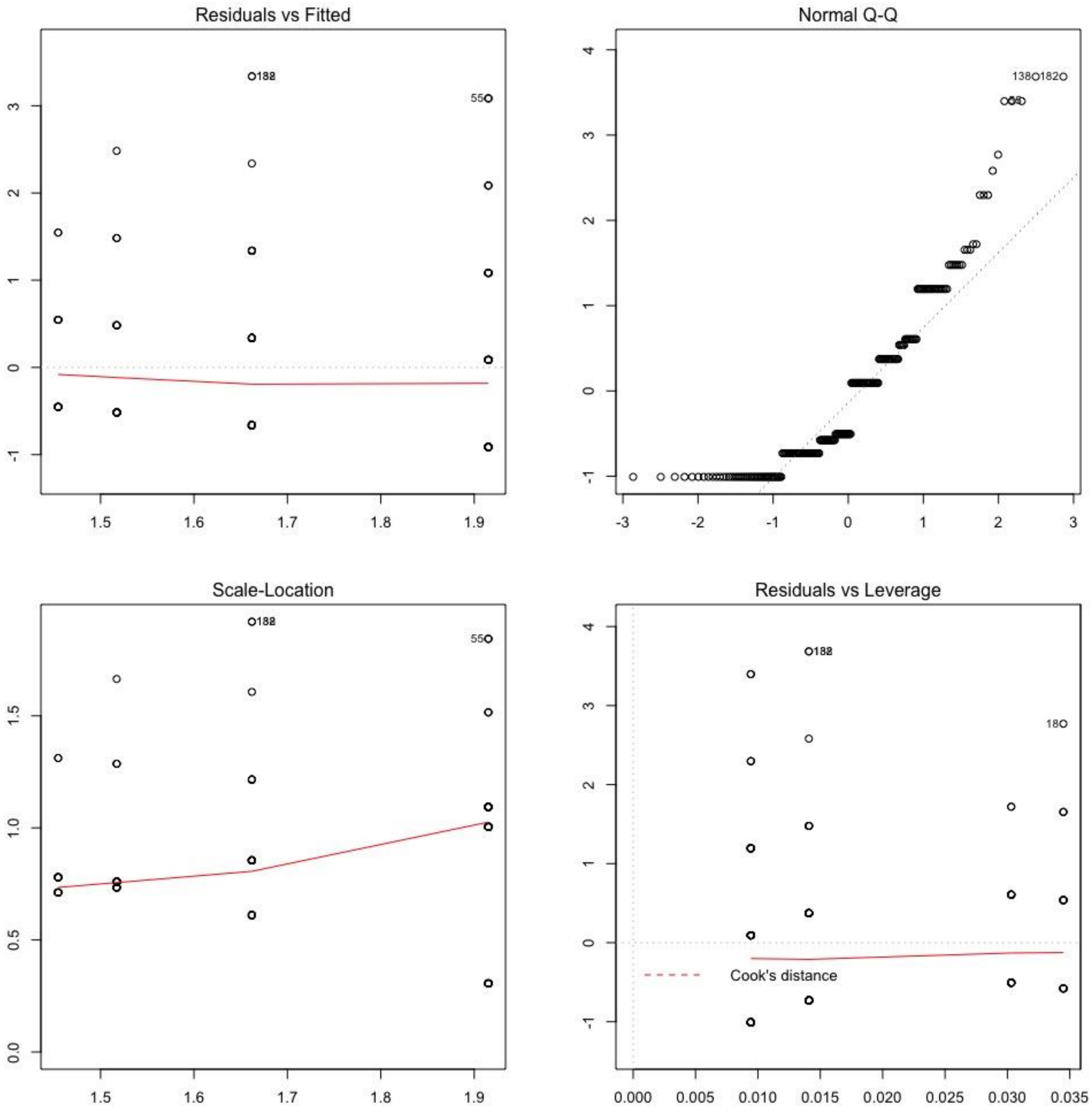


**Figure S11.** Standardised Pearson residuals from the one-way ANOVA comparing taxon richness between different eDNA isolation protocols.

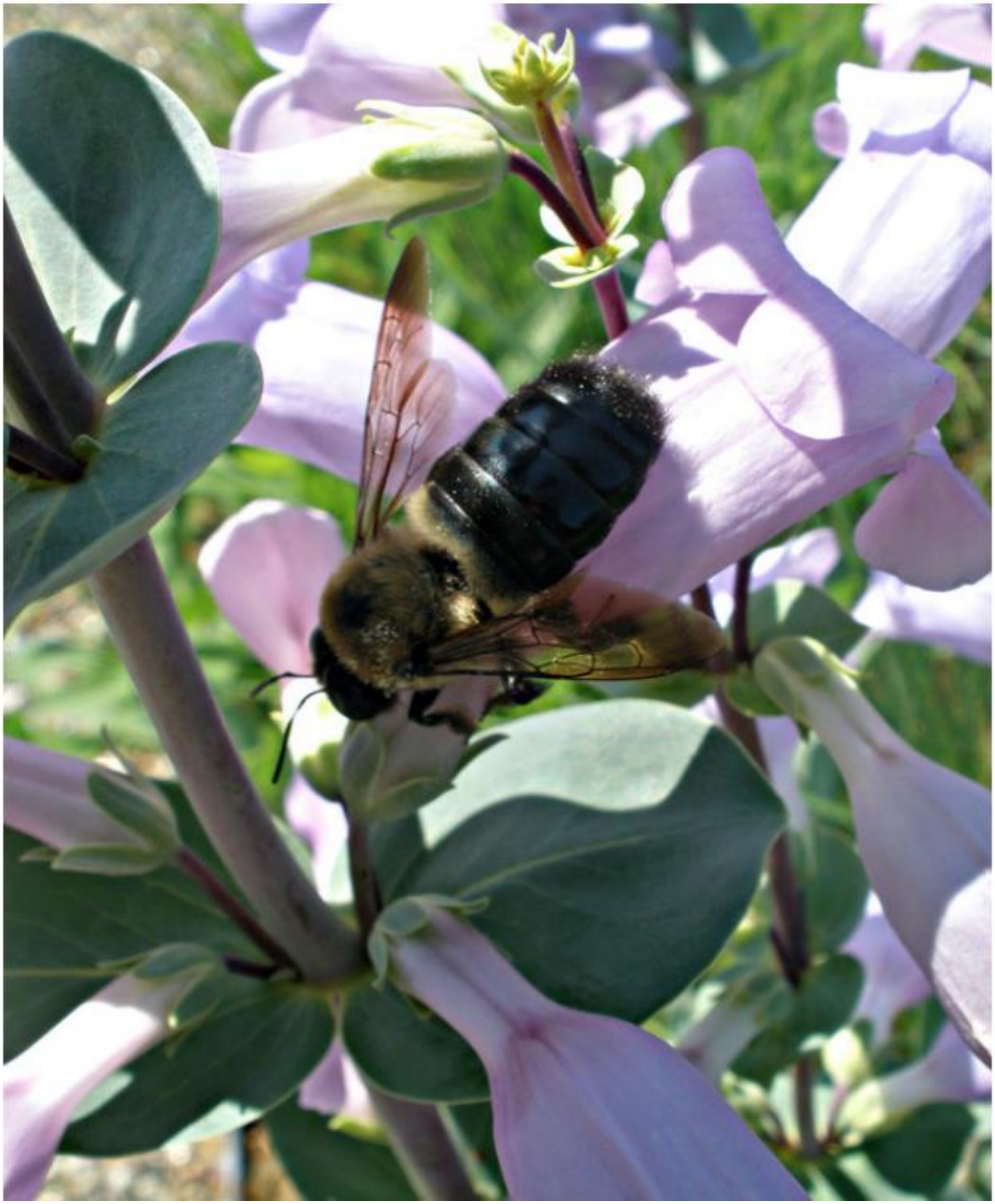

**Figure S12.** An eastern carpenter bee (*Xylocopa virginica*) exhibiting nectar robbing behavior. The bee is boring a hole by biting into the base of a *Penstemon* flower to access the nectar. Despite extensive observations and evidence of this behaviour, we failed to detect this species. Photo taken by Mark Davis.
